# Intrinsically disordered ligands for the control of receptor uptake by endocytosis

**DOI:** 10.64898/2026.04.06.716760

**Authors:** Sujeong Park, Irene Sarro, Advika Kamatar, Liping Wang, Padmini Rangamani, Eileen M. Lafer, Jeanne C. Stachowiak

## Abstract

Endocytosis plays a crucial role in signaling, recycling, and degradation of receptors. Controlling endocytosis of specific receptors is therefore a major goal for both basic science and medicine. While antibody-induced dimerization can drive signaling-induced uptake of some receptors, the steric bulk of antibodies generally inhibits endocytosis, such that control over receptor uptake remains an unmet need. Recent work has demonstrated that attractive interactions between intrinsically disordered proteins drive inward membrane curvature, a key step in endocytosis. To harness this phenomenon for the control of receptor uptake, we designed chimeras that consisted of disordered domains fused to ligands for specific receptors. These chimeras condensed on membrane surfaces, driving receptor clustering and uptake via clathrin-mediated endocytosis. In contrast, chimeras that repelled one another resisted condensation, helping receptors escape endocytosis and remain on the cell surface. Taken together, these results suggest that by modulating the amino acid sequence of intrinsically disordered ligands, we can promote or hinder the internalization of specific receptors by endocytic pathways. More broadly, these findings suggest a generalizable strategy for controlling the plasma membrane lifetime of diverse receptors, opening up new pathways for modulating cellular behavior and delivering therapeutics.

**Significance Statement:** Cells regulate signaling by continuously internalizing receptors from their surfaces using endocytosis. Controlling receptor internalization would provide new tools for addressing diverse diseases. While antibodies can cluster specific receptors on the cell surface, their steric bulk and rigidity inhibit endocytosis. In contrast, here we demonstrate that engineered ligands containing intrinsically disordered domains form flexible complexes that precisely control receptor internalization. Disordered ligands that attract one another condense receptors at sites of endocytosis, driving uptake, while repulsive disordered ligands prevent condensation such that receptors remain on the cell surface. By tuning the amino acid sequence of the disordered domain, a ligand can be switched from promoting to inhibiting receptor internalization, offering the opportunity to control cell signaling and therapeutic delivery.

## Introduction

Cell surface receptors, including receptor tyrosine kinases and G-protein coupled receptors, are central to cellular communication and signaling (1, 2). The number density of these receptors on the plasma membrane is directly linked to signalling efficiency and cellular decision making (3, 4). Cells regulate receptor abundance and tune signaling through trafficking, recycling, and degradation of receptors (5, 6). Endocytosis plays a critical role in internalizing cell surface receptors and their ligands. Eukaryotic cells utilize multiple endocytic pathways including endophilin-mediated endocytosis, macropinocytosis, and clathrin-independent pathways (7); however, the dominant endocytic route for many receptors is clathrin-mediated endocytosis (CME) (5, 8). CME proceeds in distinct stages. Adaptor proteins assemble at nascent sites of endocytosis, often coordinating with phosphatidylinositol-4,5-bisphosphate lipids (9). Subsequently, recruitment of receptors, other transmembrane proteins, and lipid cargo is coordinated with the gradual assembly of clathrin triskelia and adaptor proteins into a spherical coat (10, 11). Finally, the GTPase dynamin and its interactors oligomerize at the neck of the endocytic bud, where their conformational changes drive membrane scission, releasing cargo-loaded clathrin-coated vesicles into the cytoplasm (12).

Recruitment of receptors to sites of CME on the plasma membrane is driven by direct biochemical interactions between cytosolic sorting motifs and adaptor proteins (13). For example, the major adaptor protein of the clathrin pathway, AP2, binds simultaneously to diverse receptors, membrane lipids, and clathrin. The µ2 subunit of AP2 recognizes motifs such as YXXΦ, where Y is tyrosine, X is any amino acid, and Φ is a bulky hydrophobic residue, and dileucine motifs within the cytosolic domains of receptors (14, 15). These binding events promote recruitment of cargo into growing endocytic structures. In contrast, increasing the steric bulk of a receptor through dimerization, binding of bulky ligands, or glycosylation can generate repulsive interactions that reduce accumulation of receptors at endocytic sites, owing to the limited capacity for cargo within these structures (16). Notably, when this steric pressure becomes sufficiently large, it can even drive outward membrane protrusions with a curvature that is opposite to the inward membrane bending required for endocytosis (17, 18).

Given the critical role of receptor internalization, designing ligands that can control endocytic uptake has become an important goal for tuning cellular functions. It is well-known that antibody-induced dimerization or clustering of specific receptors, such as receptor tyrosine kinases, drives signaling events that trigger cellular uptake (19, 20). However, owing to the large size of antibodies and limited capacity of endocytic structures, cross-linking receptors with antibodies can create rigid membrane patches that stall endocytosis or favor alternate, less efficient endocytic routes (20). For this reason, antibody-based clustering is not a generalizable strategy for receptor uptake. To overcome these limitations, we need a means of locally clustering receptors on the plasma membrane without sterically or mechanically inhibiting the increase in membrane curvature required for endocytosis.

Intrinsically disordered proteins are uniquely suited to the task of forming flexible clusters on membranes, as the strength of the interactions among them can be varied with a high degree of precision from strongly attractive to weakly attractive, or even repulsive by varying their amino acid sequences (21). Further, recent work demonstrates that disordered proteins have an inherent capacity to modulate membrane curvature when bound to the membrane (22, 23). Specifically, many disordered domains contain repetitive sequences rich in charged or aromatic residues promoting weak, multivalent attractive interactions that promote liquid-liquid phase separation (LLPS) (24, 25). When disordered domains with these net attractive interactions, such as the low complexity domain of fused in sarcoma (FUSLC) or the RGG domain of LAF-1, assemble on membrane surfaces, they can generate compressive stresses that bend the membrane inward, producing protein-lined invaginations and tubules (22). In contrast, disordered domains that have net repulsive interactions with one another have been shown to drive assembly of protein-coated membrane buds and tubules in the opposite direction to those created by attractive interactions (16).

These findings raise the question of whether engineered ligands, consisting of disordered proteins, could be used to regulate receptor uptake through tunable attractive and repulsive interactions. Here we test this hypothesis by designing chimeras consisting of a ligand for a specific model receptor, conjugated with an intrinsically disordered domain. We found that chimeras that attracted one another promoted the recruitment of receptors into endocytic structures and enhanced receptor uptake. Consistent with this finding, strengthening attractive interactions by mutating the sequence of the disordered domain increased enrichment of receptors at endocytic sites, while attenuating attractive interactions decreased receptor uptake. In contrast, chimeras that repelled one another depleted receptors from endocytic sites, reducing their uptake. Taken together, these results illustrate that designing disordered protein-based chimeric ligands enable both positive and negative control over receptor internalization by endocytic pathways. This capability could be used to control cell behavior for therapeutic purposes. For example, driving rapid uptake could attenuate aberrant signaling of oncogenic receptors, while preventing internalization could sustain signaling of therapeutic targets.

## Results

### Attractive interactions between disordered ligands promote endocytosis and inward membrane curvature

We began by evaluating the impact of disordered protein ligands on membrane curvature, a key early step in endocytosis. Here we reasoned that ligands which are capable of driving inward, concave membrane curvature could bias receptors toward inwardly curved endocytic sites. The first disordered ligand we considered was the low complexity domain of Fused in Sarcoma (FUS LC), residues 1 to 163, which is enriched in serine, tyrosine, glycine, and glutamine residues (26, 27). FUS LC undergoes liquid-liquid phase separation (LLPS) through π–π stacking and dipole-dipole interactions among amino acid side chains (24). We chose FUS LC (1–163) because it is among the best studied examples of a relatively short disordered sequence that undergoes LLPS (25), and because it has been previously shown to drive inward, concave membrane bending (22). We began by evaluating this assumption, as shown in Fig. 1 *A*-*C*. Here GFP-FUS 1-163 was recruited to the surfaces of giant unilamellar vesicles (GUVs) through the interaction between its N-terminal histidine tag and membrane lipids with Ni-NTA head groups (Fig. 1*A*). When GUVs were exposed to 1 μM his-GFP-FUS 1-163, we observed protein recruitment to GUV surfaces within minutes, followed by the clear emergence of inwardly directed membrane tubules (Fig. 1 *B* and *C*). In contrast, GUVs incubated with his-GFP, without the FUS 1-163 domain, showed no increase in the incidence of membrane tubules above control samples that were not exposed to protein (Fig. 1 *B* and *C*).

**Fig. 1.**
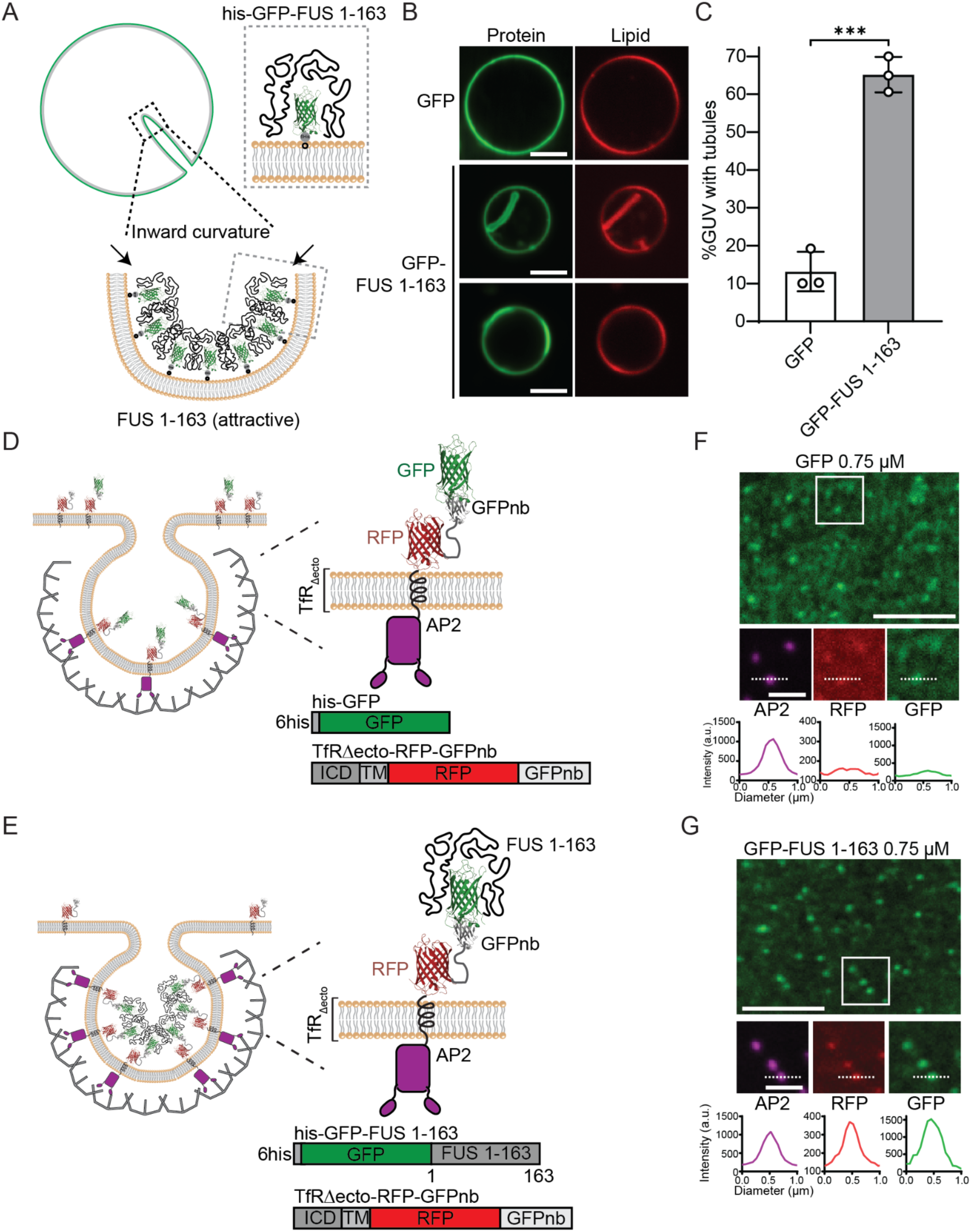
Attractive interactions between disordered ligands promote endocytosis and inward membrane curvature. **A.** Pictorial representation of GUV tubulation by his-GFP-FUS 1-163. Green lines represent FUS 1-163 protein; grey lines indicate membrane; gray dots indicate 6×histidine tags; black dots represent Ni-NTA lipids. **B.** Representative images of GUV incubated with 1 μM his-GFP (top) and his-GFP-FUS 1-163 (bottom) in 25 mM HEPES, 150 mM NaCl, pH 7.4. The scale bars are 5 µm. **C.** Fraction of GUVs exhibiting inwardly directed membrane tubules. Error bars represent mean ± SD of three independent experiments, with n > 100 GUVs in total. Statistical significance was determined by an unpaired, two-tailed Student’s *t* test. (***P < 0.001). **D-E.** Cartoon schematic illustrating endocytosis, and the transferrin receptor (TfR)-based model receptor and ligand binding. The model receptor consists of the intracellular and transmembrane domains of the TfR fused to extracellular RFP and GFP nanobody (GFPnb) domains. **(D)** GFP and **(E)** GFP-FUS 1-163 ligands bind specifically to the extracellular GFPnb domain (right). **F-G.** Spinning disk confocal microscopy images of the plasma membrane of SUM159 cells transiently expressing the model receptor. Endocytic sites were visualized using HaloTag-JF646-labeled AP2 (magenta). Cells were incubated with **(F)** 0.75 µM GFP and **(G)** 0.75 µM GFP-FUS 1-163, respectively (green). White boxes indicate the magnified insets, and graphs show intensity line profiles across the dashed lines. The scale bars are 5 µm (main) and 1 µm (insets).

Motivated by these findings, we set out to determine whether disordered ligands with attractive interactions could promote uptake of receptor-ligand complexes during cellular endocytosis. For this purpose, we made use of a chimeric transmembrane model receptor (Fig. 1 *D* and *E*). This receptor consisted of the intracellular and transmembrane domain of transferrin receptor (TfR), known for its near exclusive, constitutive internalization by clathrin-mediated endocytosis (CME) (28), fused to an extracellular red fluorescent protein (RFP) domain for visualization, followed by a single domain antibody against GFP (GFPnb), for the purpose of recruiting GFP-tagged ligand (29). To measure the uptake of the model receptor-ligand complex in the absence of disordered protein interactions, GFP alone was used as the ligand. Additional ligands were formed by fusing disordered domains to GFP, where the difference in endocytic uptake of receptor-ligand complexes with and without disordered domains was used to gauge the impact of these domains on endocytosis.

To evaluate endocytic uptake of the model receptor-ligand complex, we began by examining confocal microscopy images after transiently transfecting the model receptor into SUM159 human breast tumor epithelial cells. These cells, a gift of the T. Kirchhausen Lab, were gene-edited to add a HaloTag to the σ2 subunit of AP2, the major adaptor protein of the clathrin pathway (14, 30), enabling visualization with a HaloTag ligand bound to the JF646 fluorophore. We tracked AP2 rather than clathrin itself because AP2 localizes exclusively to the plasma membrane, whereas clathrin also associates with endosomes and other intracellular compartments (9, 31). SUM159 cells are commonly used to visualize endocytic structures because of their well-spread lamellipodia, which aid visualization of endocytic structures (32).

We observed that the receptor-ligand complex colocalized with AP2 puncta when cells were exposed to the GFP ligand (Fig. 1*F*). Interestingly, when GFP-FUS 1-163 was used as the ligand, the intensities of both GFP and RFP at AP2 puncta increased substantially compared to cells exposed to GFP alone at the same concentration (0.75 µM) (Fig. 1*G*). This enrichment of receptor-ligand complexes was notably absent when a monoclonal IgG against the model receptor was used as the ligand (*SI Appendix*, Fig. S1 *A* and *B*). Taken together, these initial observations suggested that the FUS 1-163 domain contributed to recruitment of receptor-ligand complexes to sites of clathrin-mediated endocytosis. We next sought to quantify these effects and determine their dependence on ligand concentration.

### An intermediate concentration of FUS LC ligands maximizes endocytic uptake

To quantify ligand and receptor recruitment to endocytic sites, we used CMEanalysis, a publicly available algorithm that fits a 2D Gaussian function to each puncta in the AP2 channel (JF646) to determine its intensity relative to the local background signal (33). Using these AP2 puncta as the “master” channel, the algorithm then fits 2D Gaussian functions to co-localized puncta in the “subordinate” channels corresponding to the receptor (RFP) and ligand (GFP) to determine their respective intensities. From these data, we plotted the relative intensity of the ligands at each endocytic site as a function of their intensity on the surrounding surface of the plasma membrane. The resulting plot shows a steady increase in ligand recruitment to endocytic sites with increasing ligand intensity at the cell surface (Fig. 2*A*). Notably, the slope of the rise was higher for GFP-FUS 1-163 than it was for GFP, suggesting that the FUS 1-163 domain increased recruitment of the ligand to endocytic sites, even when both ligands were added at the same concentration (0.75 µM) and present at the same relative concentration on the membrane surface (same horizontal range on plot).

**Fig. 2.**
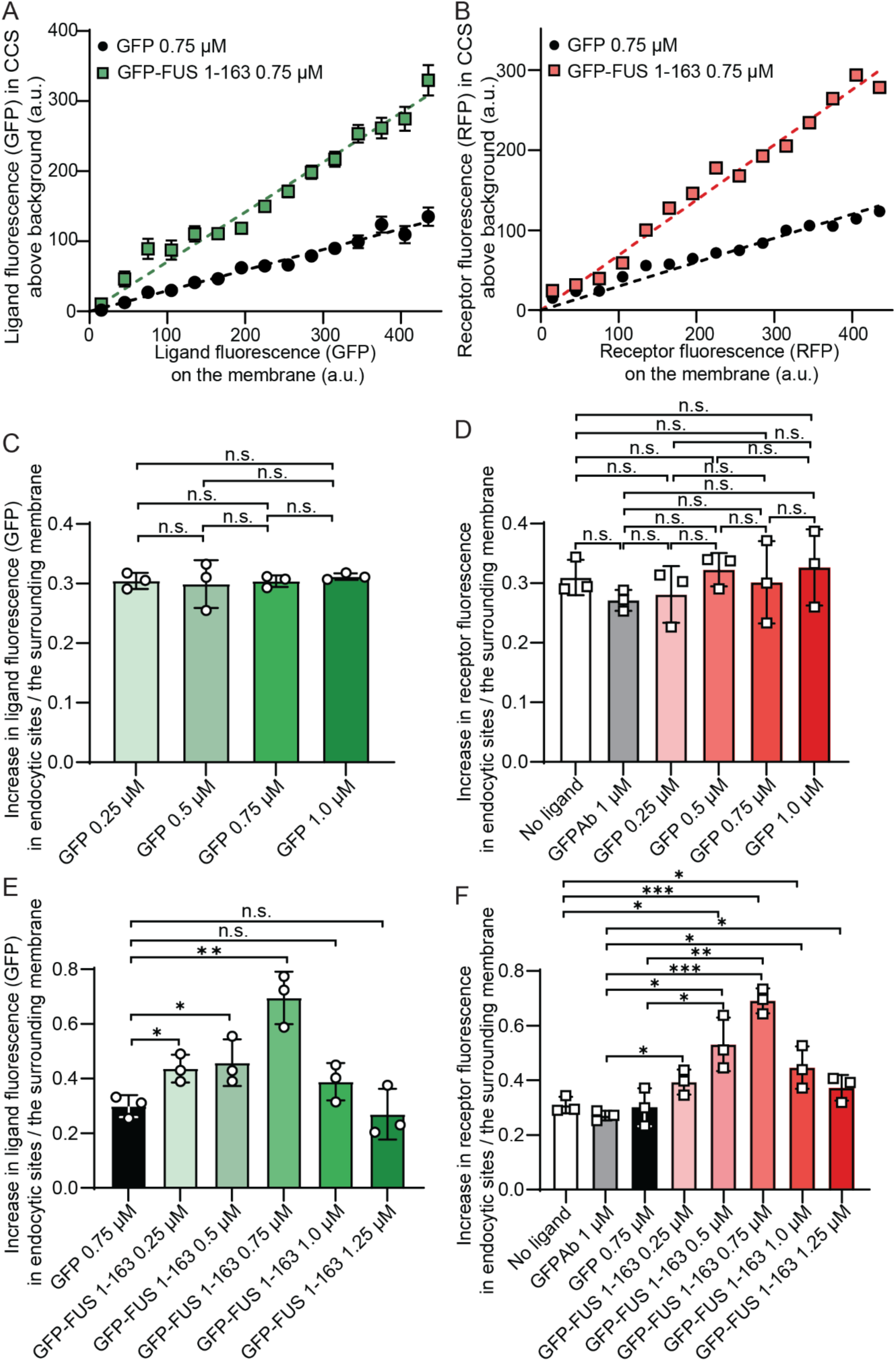
An intermediate concentration of FUS LC ligands maximizes endocytic uptake. **A-B.** The representative curves of relative number of model ligands **(A)** and receptors **(B)** within endocytic proteins localized in clathrin-coated structures is shown relative to the concentration of model ligands and receptors on the plasma membrane around each structure. A total of 4,907 endocytic structures were detected from 44 cells incubated with GFP, 10,366 endocytic structures were detected from 54 cells incubated with 0.75 µM GFP-FUS 1-163. **C-D**. Slope analysis of **(C)** ligands and **(D)** receptors for various concentrations of GFP ligands. The relative number of model receptors within endocytic proteins localized in clathrin-coated structures is shown against the relative concentration of fusion proteins on the plasma membrane around each structure. A total of 4793 endocytic structures were measured from 45 cells, 8568 endocytic structures were detected from 61 cells incubated with 1 µM GFP monoclonal antibody, 22,943 endocytic structures were detected from 148 cells incubated with 0.25 µM GFP, 40,327 endocytic structures were measured from 215 cells incubated with 0.5 µM GFP, 13,618 endocytic structures were detected from 138 cells incubated with 0.75 µM GFP, and 30,119 endocytic structures were detected from 158 cells incubated with 1.0 µM GFP. **E-F**. Plot showing slope analysis of **(E)** ligands and **(F)** receptors for various concentrations of GFP-FUS 1-163 ligands. A total of 4793 endocytic structures were measured from 45 cells, 8568 endocytic structures were detected from 61 cells incubated with 1 µM GFP monoclonal antibody, 13,618 of endocytic structures were detected from 138 cells incubated with GFP, 25,568 endocytic sites were detected from 170 cells incubated with 0.25 µM GFP-FUS 1-163, 31,340 endocytic sites were detected from 137 cells incubated with 0.5 µM GFP-FUS 1-163, 38,646 endocytic sites were detected from 155 cells incubated with 0.75 µM GFP-FUS 1-163, 27,022 endocytic sites were detected from 163 cells incubated with 1.0 µM GFP-FUS 1-163, and 36,732 endocytic sites were detected from 144 cells incubated with 1.25 µM GFP-FUS 1-163. N = 3 biologically independent samples for each group. Error bars represent mean ± SD (*P < 0.05, **P < 0.01, ***P < 0.001, n.s. means no significant difference).

We next examined whether enhanced ligand accumulation at endocytic sites correlated with enhanced recruitment of the model receptor. Using the same approach as in Fig. 2*A*, we plotted receptor intensity at endocytic sites relative to its local concentration on the plasma membrane. These data again revealed linearly increasing trends for which the slope was substantially higher in cells exposed to GFP-FUS 1-163 compared to those exposed to GFP (Fig. 2*B*). Notably, neither the fluorescence intensity of AP2 within endocytic sites, which provides a measure of their relative size, nor the density of endocytic sites on the plasma membrane varied between cells exposed to GFP versus GFP-FUS 1-163 (*SI Appendix,* Fig. S2). These data suggest that the increased intensity of receptors at endocytic sites corresponds to an increase in receptor density within these structures, rather than changes to their size or number.

Next we varied the concentration of the ligands to determine their impact on recruitment of receptor-ligand complexes to endocytic sites (0.25 µM to 1.25 µM). For each concentration, we constructed plots of ligand and receptor intensity at endocytic sites as a function of their intensity on the plasma membrane (*SI Appendix,* Fig. S3). Fitting these data with linear trends, we plotted the slopes as a function of ligand concentration (Fig. 2 *C* and *D* and *SI Appendix,* Fig. S3 *A* and *B*). These slopes were approximately constant across all ligand concentrations, as were the concentrations of ligand on the plasma membrane (*SI Appendix*, Fig. S4 *A* and *B*), suggesting that binding of the GFP ligand to cells was controlled by cellular expression of the model receptor, rather than the ligand concentration. Notably, there was no increase in enrichment of the model receptor at endocytic sites for any concentration of the GFP ligand. Similarly, a monoclonal IgG against the model receptor failed to increase its recruitment to endocytic sites, in line with literature reporting that conventional antibodies do not promote clathrin-mediated endocytosis (20).

In contrast, for cells exposed to the GFP-FUS 1-163 ligand, the slopes of the recruitment curves increased with increasing ligand concentration, reaching a maximum value at a concentration of 0.75 µM, before decreasing (Fig. 2 *E* and *F* and *SI Appendix,* Fig. S3 *C* and *D*). Meanwhile, the concentration of ligand recruited to the surrounding plasma membrane increased monotonically with increasing ligand concentration (*SI Appendix*, Fig. S4 *C* and *D*). Notably, the same trend was observed for a ligand consisting of a longer form of the FUS LC domain, GFP-FUS 1-214 (*SI Appendix*, Fig. S5 *A*-*D*) (34). Here the optimal concentration for recruitment of ligands and receptors to endocytic sites shifted slightly to 0.5 µM for GFP-FUS 1-214 (*SI Appendix*, Fig. S5 *E* and *F*), perhaps due to slightly stronger attractive interactions among the longer FUS LC domains. How can the decrease in receptor-ligand localization to endocytic sites at high ligand concentrations be explained?

### A high concentration of FUS LC ligands creates plaque-like condensates that oppose endocytic uptake of ligands and receptors

Interestingly, coincident with the reduction in recruitment of ligands and receptors to endocytic sites at increasing concentrations of GFP-FUS 1-163, we also observed the emergence of irregularly shaped, micrometer-scale plaque-like condensates on the plasma membrane, which were intense in both the ligand and receptor fluorescence images (Fig. 3*A*). While these structures did not colocalize clearly with endocytic sites in the AP2 images, endocytic sites were present on the surrounding plasma membrane, where they continued to concentrate ligands and receptors (Fig. 3*A*, insets), albeit at a lower level than observed at lower ligand concentrations (Fig. 2).

**Fig. 3.**
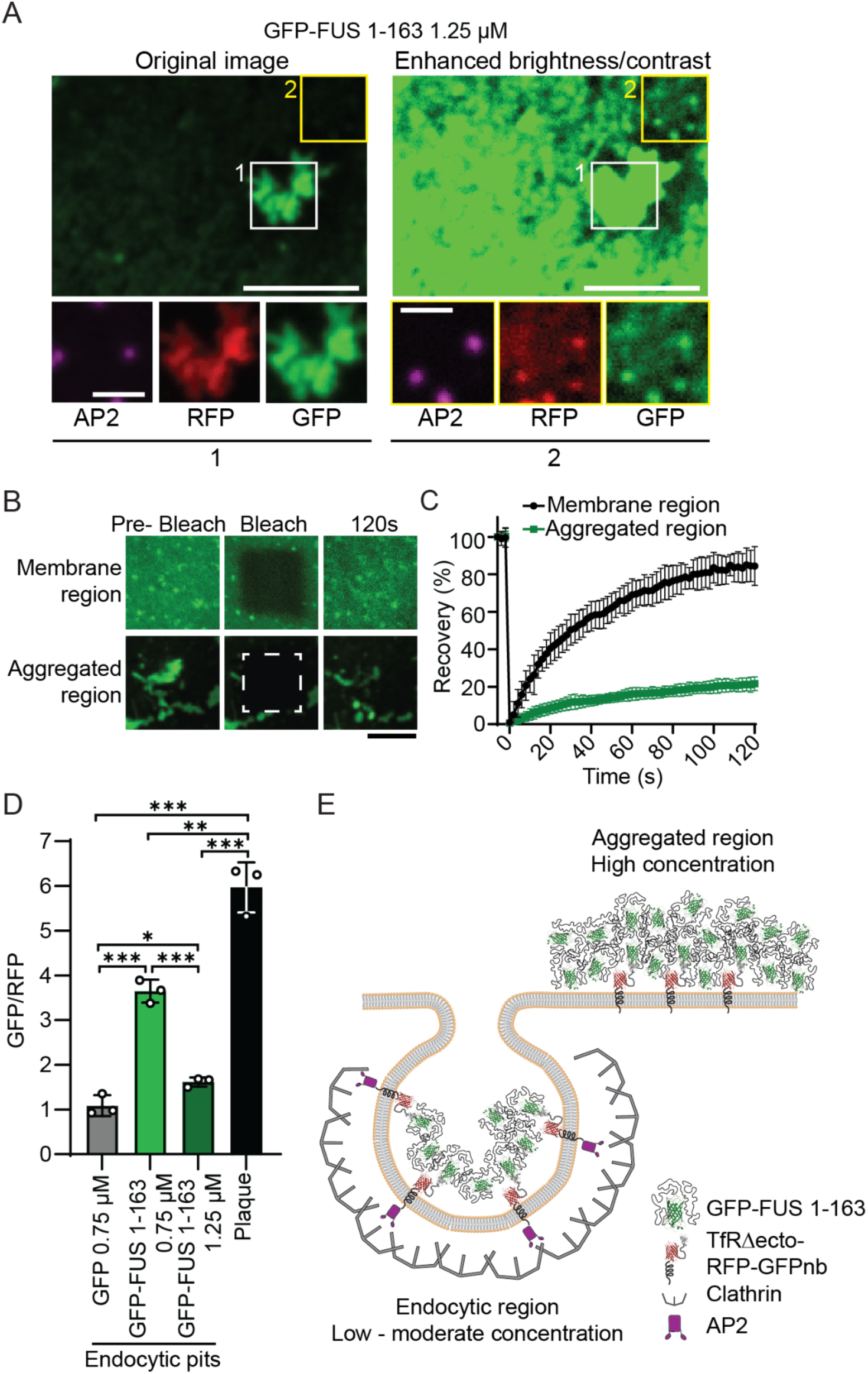
A high concentration of FUS 1-163 ligands creates plaque-like structures that oppose endocytic uptake of ligands and receptors. **A.** Spinning disk confocal microscopy images of the plasma membrane of SUM159 cells transiently expressing the model receptor. Endocytic sites were visualized using HaloTag-JF646-labeled AP2 (magenta). Cells were incubated with 1.25 µM GFP-FUS 1-163 (green). Left: original images showing aggregated regions on the plasma membrane and insets of region 1. Right: the same fields with enhanced brightness/contrast to reveal dim puncta in region 2. White and yellow boxes indicate the magnified insets. The scale bars are 5 µm (main) and 1 µm (insets). **B.** Confocal images of fluorescence recovery after photobleaching (FRAP) at the plasma membrane of SUM159 cells incubated with 1.25 µM GFP-FUS 1-163. A square region of 5 µm on each side was bleached. The dashed square indicates a bleached region. The scale bar is 5 µm. **C.** Fluorescence recovery curves for membrane and aggregated regions. Lines are average fluorescence recovery ± SD. N = 6 regions were analyzed for each plot. **D**. Bar graph shows the ratio of the approximate number of ligands (GFP) to receptors (RFP), estimated using single-molecule calibration (*SI Appendix*, Fig. S6). Data compares endocytic pits in cells treated with 0.75 µM GFP, 0.75 µM GFP-FUS 1-163, and 1.25 µM GFP-FUS 1-163, alongside aggregated plaques formed at 1.25 µM. N = 3 biologically independent samples for each group. Error bars represent mean ± SD (*P < 0.05, ***P < 0.001, n.s. means no significant difference). **E.** Schematic showing how GFP-FUS 1-163 behaves differently at low - moderate versus high concentrations. At low - moderate concentrations, GFP-FUS 1-163 remains dynamic and can accumulate within endocytic sites. At high concentrations, GFP-FUS 1-163 forms surface aggregates, reducing its availability for recruitment to endocytic sites.

To probe the assembly state of these plaque-like condensates, we performed fluorescence recovery after photobleaching (FRAP) experiments (Fig. 3*B*). In the membrane region surrounding the plaques, GFP-FUS 1-163 fluorescence recovered almost completely after photobleaching (84.39 ± 10.39%), consistent with free diffusion of receptors and ligands on the plasma membrane. In contrast, recovery was greatly reduced in the plaque-like condensates (21.65 ± 3.69%), consistent with a strongly assembled state with lower molecular turnover (Fig. 3*C*).

Given that plaque-like condensates emerged at higher concentrations of the disordered ligand, they may contain a higher ligand to receptor ratio than other areas of the plasma membrane. Therefore, we next sought to determine the ligand to receptor ratio within plaques, endocytic sites, and the surrounding plasma membrane. To estimate the ligand to receptor ratio in our experiments, we performed single-molecule fluorescence calibration experiments. These experiments measured the absolute fluorescence intensity of GFP and RFP under our imaging conditions (*SI Appendix*, Fig. S6). Using the ratio of these values, we were able to convert the ratio of fluorescence intensities for ligands and receptors to approximate stoichiometric ratios (Fig. 3*D*).

For the control GFP ligand, the ratio of ligand to receptor within endocytic structures was approximately 1:1, as would be expected for simple biochemical binding between receptors and ligands without ligand-ligand interactions. In contrast, for GFP-FUS 1-163 at the optimal concentration (0.75 µM), the GFP to RFP ratio significantly increased to 3-4:1, suggesting that attractive interactions between the ligands facilitated the condensation of multiple ligands per receptor. However, at higher concentrations (1.25 µM), the ligand to receptor ratio within endocytic structures sites decreased to around 1-2:1 (Fig. 3*D* and *SI Appendix,* S7 *A* and *B*), despite a monotonic increase in the ratio of ligands to receptors on the surrounding plasma membrane (*SI Appendix*, Fig. S7 *C* and *D*). Meanwhile, the ligand to receptor ratio in plaque-like condensates was approximately 6:1, substantially higher than in endocytic sites, even at the optimum concentration. Collectively, these results suggest that as the concentration of disordered ligands increased, ligand-ligand interactions led to condensation of ligands and receptors at endocytic sites. At moderate ligand concentrations, this condensation helped to concentrate receptors at endocytic sites, promoting their internalization. However, when the ligand concentration became too high, larger condensates with a higher ratio of ligands to receptors formed on the plasma membrane, where they sequestered both receptors and ligands away from endocytic sites, suppressing receptor internalization (Fig. 3*D*). These findings highlight the potential importance of tuning the relative strength of ligand-receptor and ligand-ligand interactions to optimize the ability of disordered protein condensates to promote endocytic uptake of receptors.

### The strength of attractive interactions among ligands controls the recruitment of receptors to endocytic structures

We next asked to what extent it is possible to modulate endocytic uptake of receptors by varying the strength of attractive interactions among disordered ligands. Specifically, we made mutations to the FUS 1-163 domain that are known to either strengthen or weaken its condensation. In particular, we targeted glutamine residues, which are critical for phase separation through hydrogen bonding (35). Previous studies have shown that increasing glutamine content enhances both phase separation and inward membrane bending (22, 25). Therefore, we utilized previously reported mutants in which either 8 glutamine residues were mutated to serines (Q to S), or 12 serine residues were mutated to glutamine (S to Q) (Fig. 4 *A*-*C*). We first acquired confocal fluorescent images of cells that expressed the model receptor and were exposed to one of the three ligands: GFP-FUS 1-163, GFP-FUS 1-163 QtoS or GFP-FUS 1-163 StoQ. All three ligands were recruited to the plasma membrane of cells expressing the model receptor. Compared to GFP-FUS 1-163, the QtoS mutant exhibited weaker colocalization with endocytic sites, while the StoQ mutant showed stronger colocalization (Fig. 4 *D*-*F*).

**Fig. 4.**
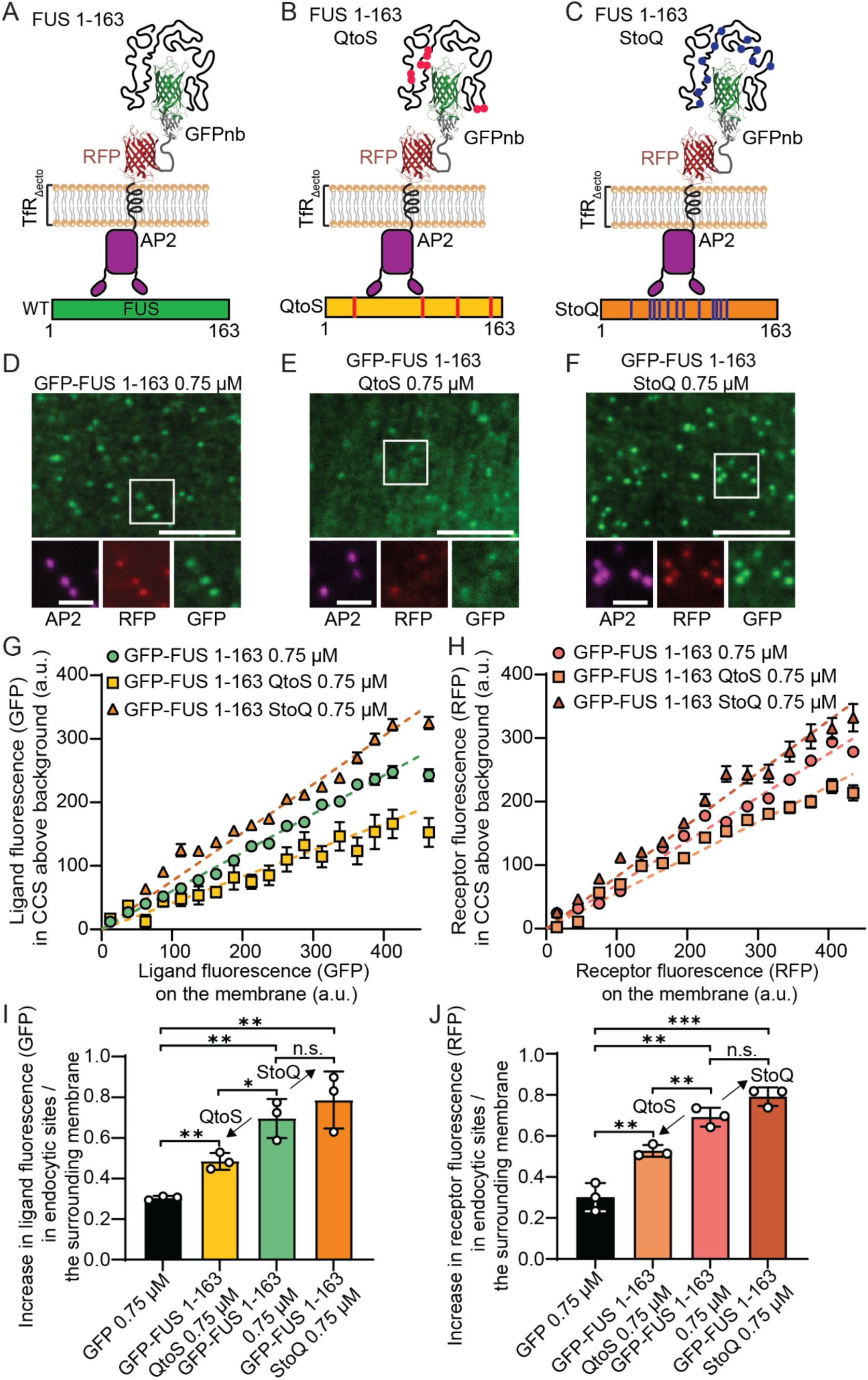
Strength of attractive interactions among ligands Degree of phase separation controls the recruitment of ligands and receptors to endocytic structures A-C. A schematic of the model receptors bound to **(A)** wild-type GFP-FUS 1-163, **(B)** GFP-FUS 1-163 QtoS, in which 8 glutamine residues were replaced with serines, and **(C)** GFP-FUS 1-163 StoQ, in which 12 serines were mutated to glutamines. **D-F.** Spinning disk confocal microscopy images of the plasma membrane of SUM159 cells transiently expressing the model receptor, visualized with HaloTag-JF646-labeled AP2 (magenta). Cells were incubated with 0.75 µM of **(D)** GFP-FUS 1-163, **(E)** GFP-FUS 1-163 QtoS, or **(F)** GFP-FUS 1-163 StoQ (green). The white boxes indicate the magnified insets. The scale bars are 5 µm (main) and 1 µm (insets). **G-H.** The representative relative number of model **(G)** ligands and **(H)** receptors within endocytic proteins localized in endocytic structures is shown against the relative concentration of fusion proteins on the plasma membrane around each structure. Error bars represent mean ± SE. A total of 10,366 endocytic sites were detected from 54 cells incubated with 0.75 µM GFP-FUS 1-163, 6,713 endocytic sites were detected from 49 cells incubated with 0.75 µM GFP-FUS 1-163 QtoS, and 6,279 endocytic sites were measured from 46 cells incubated with 0.75 µM GFP-FUS 1-163 StoQ. **I-J.** Slope analysis of increase in **(I)** ligand and **(J)** receptor fluorescence in endocytic sites versus the surrounding membrane for the three ligands. A total of 13,618 endocytic sites were detected from 138 cells incubated with 0.75 µM GFP, 38,646 endocytic sites were detected from 155 cells incubated with 0.75 µM GFP-FUS 1-163, 28,761 endocytic sites were detected from 157 cells incubated with 0.75 µM GFP-FUS 1-163 QtoS, and 42,756 endocytic sites were measured from 158 cells incubated with 0.75 µM GFP-FUS 1-163 StoQ. N = 3 biologically independent samples for each group. Error bars represent mean ± SD from three independent experiments. Statistical significance was tested using an unpaired, two-tailed Student’s *t* test (*P < 0.05, **P < 0.01, ***P<0.001, n.s. means no significant difference).

To quantify these differences, we constructed recruitment curves similar to those in Fig. 2*D*. We measured both the ligand and receptor intensity within endocytic structures and their intensity on the surrounding membrane. From these plots, we observed that ligand and receptor recruitment to endocytic sites increased with increasing interaction strength among the ligands (Fig. 4 *G* and *H*). In particular, the slope of recruitment curve, which reflects relative recruitment of receptor-ligand complexes to endocytic sites, was decreased for cells exposed to the QtoS mutant and increased for those exposed to the StoQ mutant, relative to those exposed to GFP-FUS 1-163 (Fig. 4 *I* and *J*), resulting in a maximum receptor uptake that was more than 2.5 times greater than for the control ligand (StoQ mutant versus GFP alone). Taken together, these data suggest that modulating the strength of interaction between disordered ligands modulates the extent of endocytic uptake of the resulting receptor-ligand complexes.

### Attractive interactions among disordered ligands of diverse amino acid composition have a similar impact on receptor uptake

Next we asked to what extent our findings with ligands containing FUS LC could be generalized to other intrinsically disordered ligands that attract one another. For this purpose, we designed an additional ligand where the FUS LC domain was replaced with another well-studied disordered domain that undergoes phase separation, the RGG domain of LAF-1. The N-terminal 168 amino acids of LAF-1 are intrinsically disordered and enriched in glycine and arginine residues, including several RGG motifs (36). While attractive interactions between FUS LC domains are driven primarily by π-π interactions among aromatic residues and dipole-dipole interactions, the attractive interactions among RGG domains are driven by electrostatic interactions between positively charged arginine residues and negatively charged aspartic acid residues, along with cation-π interactions between positively charged and aromatic residues, and dipole-dipole interactions (24). Despite these differences, both domains are able to form condensates when recruited to membrane surfaces (22), suggesting the potential for shared behaviors in the context of endocytosis.

To test this idea, we fused his-GFP to the RGG domain of LAF-1. When his-GFP-RGG ligands bound to the surfaces of GUVs via histidine-nickel interactions, we observed inwardly directed membrane tubules (Fig. 5*A*), similar to those observed with his-GFP-FUS 1-163 (Fig. 1*A*). Quantification showed approximately a 4 fold increase in the percentage of vesicles with tubules for vesicles exposed to his-GFP-RGG compared to those exposed to his-GFP (Fig. 5*B*). Next, we used the model receptor system defined above to evaluate the impact of GFP-RGG on endocytic uptake of the model receptor (Fig. 5*C*). At relatively low concentration (0.125 µM), GFP-RGG colocalized clearly with endocytic sites in cells expressing the model receptor, similar to our observations with cells exposed to GFP-FUS 1-163 ligands (Fig. 5*D*). However, at higher concentration (0.5 µM), the GFP-RGG ligand began to form plaque-like condensates on the plasma membrane of exposed cells, suggesting excessive condensation of the ligand with the model receptor (Fig. 5*E*). Quantification of these data revealed that addition of 0.125 µM of GFP-RGG increased the partitioning of ligands and receptors to endocytic sites relative to cells exposed to the GFP ligand (0.75 µM) (Fig. 5 *F* and *G*). However, the magnitude of this increase declined as the concentration of GFP-RGG increased to 0.5 µM, correlating with the appearance of plaque-like condensates. After examining a range of GFP-RGG concentrations, a trend similar to what we observed with GFP-FUS 1-163 emerged, in which partitioning of ligands and receptors initially increased with increasing ligand concentration but then began to decrease as the ligand concentration was further increased (Fig. 5 *H* and *I*). Taken together with the results for the GFP-FUS 1-163 ligand, these results suggest a phenomenon in which disordered ligands that have attractive interactions with one another can condense at endocytic sites, increasing endocytic uptake of receptors. However, excessive binding of ligands to the cell surface ultimately leads to assembly of plaque-like condensates that resist endocytosis and compete with endocytic sites for receptors and ligands. Given that FUS 1-163 and RGG have substantially different amino acid sequences, the similar effect of the two ligands on receptor uptake suggests that this phenomenon may be general.

**Fig. 5.**
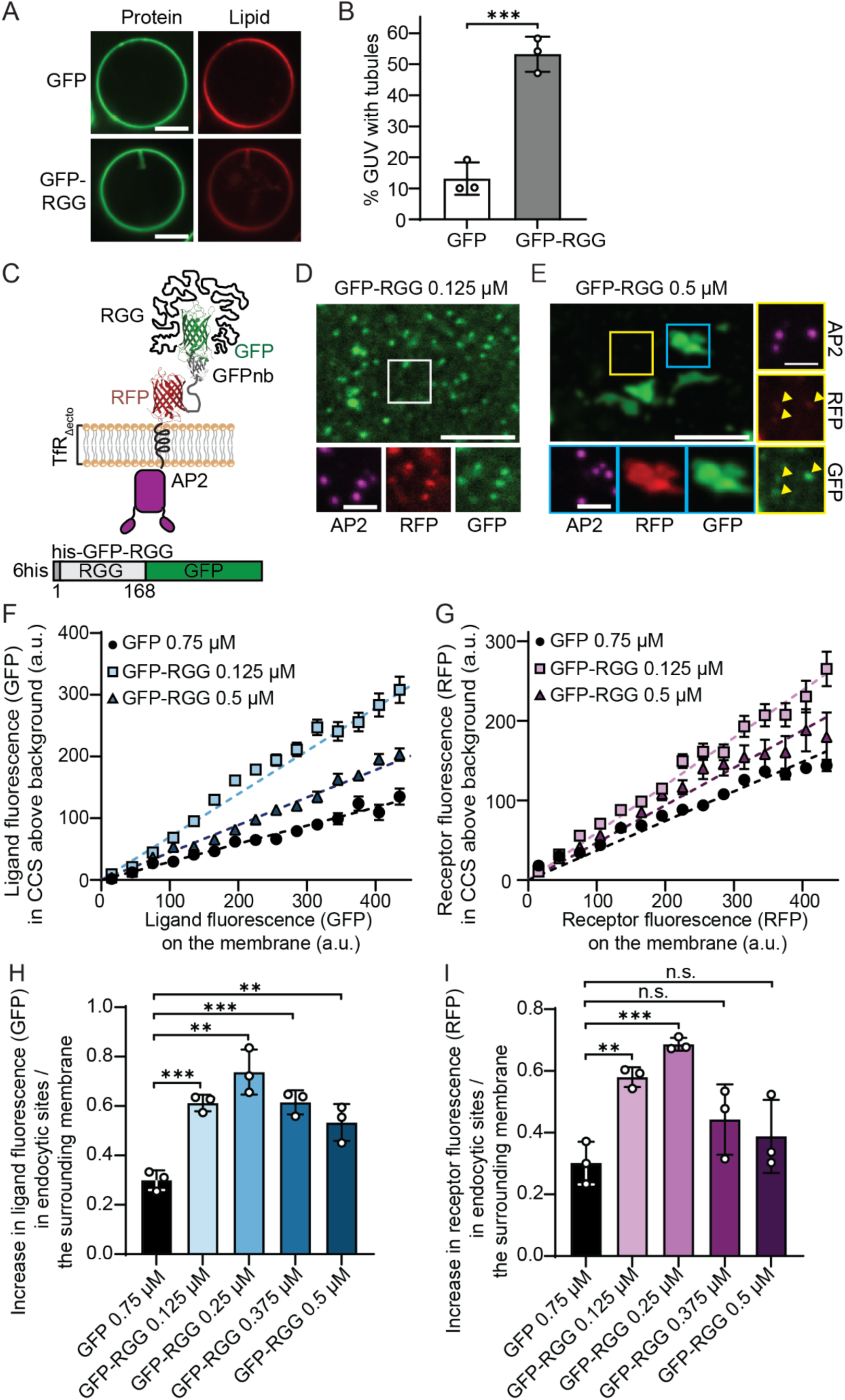
Attractive interactions among disordered ligands of diverse amino acid composition have a similar impact on receptor uptake. **A.** Representative super-resolution images of GUV incubated with 1 μM his-GFP (top) and his-GFP-RGG (bottom) in 25 mM HEPES, 150 mM NaCl, pH 7.4. The scale bars are 5 µm. **B**. Fraction of GUVs exhibiting inward curvature. Error bars represent mean ± SD of three independent experiments. n > 100 GUVs were imaged in total. Statistical significance was determined by an unpaired, two-tailed Student’s *t* test (**P < 0.01). **C.** A schematic of the model receptors bound to GFP-RGG. **D-E.** Spinning disk confocal microscopy images of the plasma membrane of SUM159 cells transiently expressing the model receptor (red), visualized with HaloTag-JF646-labeled AP2 (magenta). Cells were incubated with **(D)** 0.125 µM and **(E)** 0.5 µM of GFP-RGG. The white boxes indicate the magnified insets. The scale bars are 5 µm (main) and 1 µm (insets). **F-G.** The relative number of model **(F)** ligands and **(G)** receptors within endocytic proteins localized in endocytic structures is shown against the relative concentration of fusion proteins on the plasma membrane around each structure. A total of 4,907 endocytic sites were detected from 44 cells incubated with 0.75 µM GFP, 7,563 endocytic sites were detected from 48 cells incubated with 0.125 µM GFP-RGG, and 4,569 endocytic sites were measured from 59 cells incubated with 0.5 µM GFP-RGG. Error bars represent mean ± S.E. **H-I.** Slope analysis of **(H)** ligands and **(I)** receptors for various concentrations of GFP-RGG. A total of 13,618 endocytic sites were detected from 138 cells incubated with 0.75 µM GFP, 18,104 endocytic sites were detected from 147 cells incubated with 0.125 µM GFP-RGG, 24,218 endocytic sites were measured from 144 cells incubated with 0.25 µM GFP-RGG, 19,023 endocytic sites were detected from 137 cells incubated with 0.375 µM GFP-RGG, and 11,903 endocytic sites were detected from 163 cells incubated with 0.5 µM GFP-RGG. Error bars represent mean ± SD from three independent experiments. Statistical significance was tested using an unpaired, two-tailed Student’s *t* test (**P < 0.01, ***P < 0.001, n.s. means no significant difference).

### Mutation of RGG to reduce attractive interactions between ligands results in reduced endocytic uptake of ligands and receptors

Next we asked whether weakening the biochemical interactions responsible for association between RGG domains within ligands would weaken receptor uptake, similar to our findings with ligands containing FUS 1-163 (Fig. 4). For this purpose, we used a mutant of the LAF-1 RGG domain in which all 23 arginine residues were substituted with lysine residues, GFP-KGG (37). This mutant preserves the net charge of the RGG domain but weakens the cation–π interactions important for phase separation, owing to the inferior ability of lysine to form such interactions in comparison to arginine (37). Consistent with reduced attraction between the ligands, GUVs exposed to 1 µM his-GFP-KGG showed outwardly directed membrane curvature, in contrast to inward tubules formed by his-GFP-RGG (Fig. 6*A*). Quantification confirmed a strong decrease in the fraction of vesicles displaying inward tubules, together with an increase in outward tubules for vesicles exposed to his-GFP-KGG (Fig. 6*B*). Interestingly, these results suggest that the interactions between GFP-KGG proteins include both repulsive and attractive components, rather than being strongly attractive, as was the case for GFP-RGG.

**Fig. 6.**
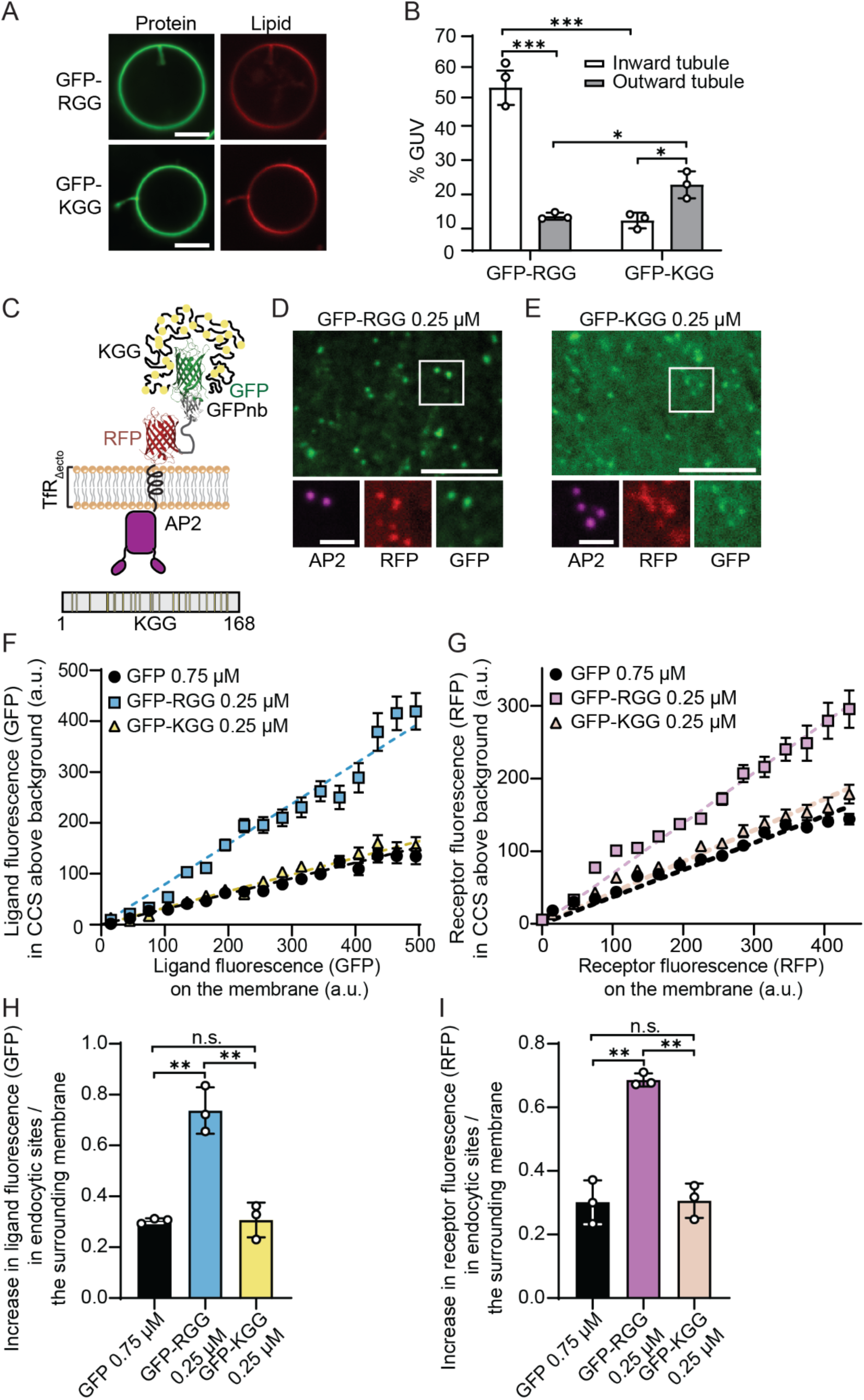
Mutation of RGG to reduce attractive interactions between ligands results in reduced endocytic uptake of ligands and receptors. **A.** Representative super-resolution images of GUV incubated with 1 μM his-GFP-RGG (top) and his-GFP-KGG (bottom) in 25 mM HEPES, 150 mM NaCl, pH 7.4. The scale bars are 5 µm. **B.** Fraction of GUVs exhibiting inward and outward curvature. Error bars represent mean ± SD of three independent experiments, with n > 100 GUVs in total. Statistical significance was determined by an unpaired, two-tailed Student’s *t* test. (*P < 0.05, ***P < 0.001). **C.** Schematic of illustration of TfR model receptors bound to GFP-KGG ligands. In the GFP-KGG ligand, all 23 arginine residues in wild-type RGG were replaced with lysine to disrupt phase separation behavior. The receptor consists of the intracellular and transmembrane domain of the TfR fused to an extracellular RFP and GFPnb. **D-E.** Spinning disk confocal microscopy images of the plasma membrane of SUM159 cells transiently expressing the model receptor, visualized with HaloTag-JF646 conjugated AP2 (magenta). Receptors are incubated with **(D)** 0.25 µM GFP-RGG or **(E)** 0.25 µM GFP-KGG. The white boxes indicate the magnified insets. The scale bars are 5 µm (main panels) and 1 µm (insets). **F.** The relative number of model **(F)** ligands and **(G)** receptors within endocytic proteins localized in endocytic structures is shown against the relative concentration of fusion proteins on the plasma membrane around each structure. A total of 4,907 endocytic structures were detected from 44 cells incubated with 0.75 µM GFP, 7,190 endocytic sites were detected from 43 cells incubated with 0.25 µM GFP-RGG, 9,584 endocytic structures were detected from 44 cells incubated with 0.25 µM GFP-KGG. Error bars represent mean ± S.E. **H-I.** Slope analysis of **(H)** ligands and **(I)** receptors comparing 0.75 µM GFP, 0.25 µM GFP-RGG and 0.25 µM GFP-KGG affinity for endocytic sites. A total of 13,618 endocytic structures were detected from 138 cells incubated with GFP, 24,218 endocytic sites were detected from 144 cells incubated with 0.25 µM GFP-RGG, 40,697 endocytic structures were detected from 152 cells incubated with 0.25 µM GFP-KGG. The individual data points represent independent samples (N = 3). Error bars represent mean ± SD. Statistical significance was tested using an unpaired, two-tailed Student’s *t* test (**P < 0.01, n.s. means no significant difference).

We next asked whether this change in curvature was accompanied by reduced recruitment of ligands and receptors to endocytic sites in cells. At 0.25 µM, GFP-RGG showed clear enrichment at endocytic sites, whereas GFP-KGG displayed visibly weaker enrichment at the same concentration (Fig. 6 *D* and *E*). To quantify these effects, we plotted recruitment curves comparing the intensity of ligands and receptors within endocytic sites to their respective intensities on the surrounding membrane for cells exposed to both ligands (Fig. 6 *F* and *G*). The slopes of these curves, which indicate the strength of partitioning of ligands and receptors to endocytic sites, were substantially reduced for cells exposed to GFP-KGG in comparison to cells exposed to GFP-RGG (Fig. 6 *H* and *I*). Similarly, these results demonstrate that weakening the affinity between RGG domains via R to K mutations results in weaker recruitment of receptors and ligands to endocytic sites. These results are similar to our findings in Fig. 4, where weakening the affinity between FUS 1-163 ligands, via the S to Q mutation, reduced recruitment of ligands and receptors to endocytic sites. The similarity between these trends further suggests that the strength of attractive interactions between disordered ligands modulates their effect on endocytosis, regardless of the specific amino acid-level interactions responsible for the attraction.

### Repulsive interactions between disordered ligands reduce endocytic uptake of ligands and receptors

Our results so far suggest that attractive interactions among disordered protein ligands facilitate endocytic uptake of receptors by promoting membrane bending and receptor clustering. However, not all disordered proteins have attractive interactions toward one another. Some exhibit net repulsive interactions. For example, disordered sequences found within endocytic adaptor proteins such as the C-terminal domains of AP180 and Epsin, which have been reported to repel one another, resulting in the ability to drive outward membrane curvature (38). Harnessing these repulsive interactions to retard endocytic uptake of receptors could be desirable as a means of prolonging cell signaling or retaining under expressed receptors on the cell surface.

To test how repulsive interactions among intrinsically disordered ligands might influence uptake of our model receptor, we replaced the attractive domains studied above (FUS LC, RGG) with a portion of the AP180 C-terminal domain (residues 328 to 518), to create GFP-AP180CTD (Fig. 7*A*) (23). This portion of AP180CTD has a net negative charge of −21, which generates strong electrostatic repulsion repulsive interactions that opposes self-association and molecular crowding (39). We first tested the effect of his-GFP-AP180CTD on the curvature of GUVs, using binding interactions between Ni-NTA lipid headgroups to attract the histidine-tagged ligand to the membrane surface. Unlike the his-GFP control, which had a minimal impact on membrane curvature, exposure of vesicles to his-GFP-AP180CTD induced assembly of mainly outwardly directed membrane tubules (Fig. 7 *B* and *C*). This result confirms that repulsive interactions among AP180CTD domains were sufficient to generate outward membrane curvature.

**Fig. 7.**
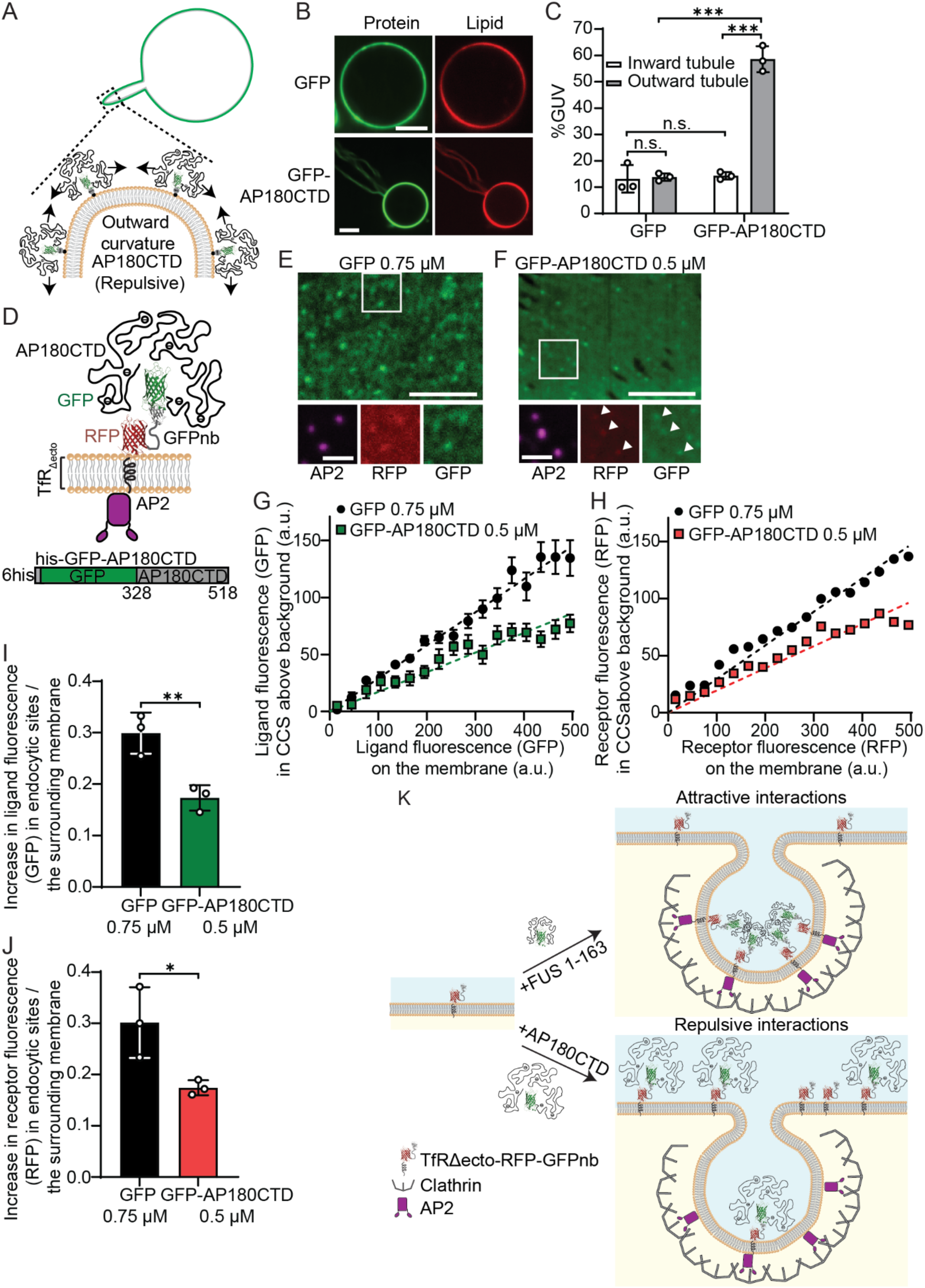
Repulsive interactions between disordered ligands reduce endocytic uptake of ligands and receptors. **A.** Pictorial representation of his-AP180CTD inward tubule formation on a GUV. Green lines represent his-AP180CTD protein, Gray lines indicate membrane, Gray dots indicate 6×histidine tags, and black dots represent Ni-NTA lipids. **B.** Representative super-resolution images of GUV incubated with 1 μM his-GFP (top) and his-GFP-AP180CTD (bottom) in 25 mM HEPES, 150 mM NaCl, pH 7.4. The scale bars are 5 µm. **C.** Fractions of GUVs exhibiting inward and outward curvature. Error bars represent mean ± SD of three independent experiments, with n > 100 GUVs in total. Statistical significance was determined by an unpaired, two-tailed Student’s *t* test. (***P < 0.001, n.s. means no significant difference). **D.** A schematic of the model receptors bound to GFP-AP180CTD. **E-F.** Spinning disk confocal microscopy images of the plasma membrane of SUM159 cells transiently expressing the model receptor (RFP), visualized with HaloTag-JF646 conjugated AP2 (magenta). Receptors are incubated with **(E)** 0.75 µM GFP or **(F)** 0.5 µM GFP-AP180CTD. The white boxes indicate the magnified insets. The scale bars are 5 µm (main panels) and 1 µm (insets). **G-H.** The relative number of model **(G)** ligands and **(H)** receptors within endocytic proteins localized in endocytic structures is shown against the relative concentration of model ligands and receptors on the plasma membrane around each structure. A total of 4,907 endocytic structures were detected from 44 cells incubated with 0.75 µM GFP, and 7,749 endocytic structures were detected from 45 cells incubated with GFP-AP180CTD. Error bars represent mean ± SE. **I-J.** Slope analysis of **(I)** ligands and **(J)** receptors affinity for endocytic sites. The individual data points represent independent samples from three trials. A total of 40,327 endocytic structures were detected from 215 cells incubated with 0.5 µM GFP, and 20,942 endocytic structures were detected from 140 cells incubated with GFP-AP180CTD. Error bars represent mean ± SD. Statistical significance was tested using an unpaired, two-tailed Student’s *t* test (**P < 0.01, *P<0.05). **K.** Schematic illustrating the opposing effects of attractive and repulsive disordered ligands on membrane curvature. Attractive phase-separable ligands such as FUS 1-163 or RGG promote inward bending, whereas repulsive ligands such as AP180CTD drive receptors out of inwardly curved membrane structures, owing to their preference for outward curvature.

Next, we examined the impact of the GFP-AP180CTD ligand on recruitment of receptors and ligands to endocytic sites in cells expressing the same model receptor used above (Fig. 7*D*). As shown in Fig. 7 *E*-*F*, GFP-AP180CTD displayed visibly reduced colocalization with endocytic sites relative to the control ligand, GFP. Notably, this trend of reduced recruitment of the disordered ligand is in clear contrast to the disordered ligands with attractive interactions, which each displayed increased recruitment to endocytic sites. Recruitment curves for ligands and receptors in Fig. 7 *G*-*H* show a downward shift for cells exposed to the GFP-AP180CTD ligand, indicating impaired ligand and receptor enrichment at endocytic sites compared to the GFP control ligand. Analysis of the slopes of these trends further confirmed that the GFP-AP180CTD ligand effectively reduced partitioning of ligands and receptors to endocytic sites (Fig. 7 *I* and *J*). Taken together, these results suggest that when disordered protein ligands have net repulsive interactions with one another, they suppress endocytic uptake of the model receptor. This reduced uptake is likely due to their tendency to induce outward membrane bending, which would be expected to mechanically oppose the inward curvature required for endocytosis (Fig. 7*K*). Overall, engineered disordered protein ligands with either attractive or repulsive interactions exerted either positive and negative control, respectively, over receptor internalization, with a dynamic range of about 4 to 5-fold between the strongest attractive ligand (GFP-FUS StoQ) and the repulsive ligand (GFP-AP180CTD).

## Discussion

Here we show that intrinsically disordered ligands can be used to modulate the endocytic uptake of cell surface receptors. While classical models of endocytosis emphasize biochemical recognition between cargo motifs and adaptor proteins (14, 15), our results highlight that the extent of biophysical attraction or repulsion between the ligands can strongly modulate the recruitment of receptors to endocytic sites. Specifically, we found that intrinsically disordered ligands with net attractive interactions, which would be expected to drive condensation *in vitro* (22), drove increased partitioning of receptors into endocytic structures. In contrast, disordered ligands with net repulsive interactions drove exclusion of receptors, likely owing to a combination of electrostatic repulsion and steric exclusion (16, 38, 40).

Our findings regarding attractive ligands, such as FUS 1-163 and RGG, align with the emerging consensus that protein condensation can drive membrane remodeling (22, 23, 41). Specifically, previous work has suggested that the condensation of disordered domains on membrane surfaces generates compressive stress, which creates spontaneous inward curvature (22). Here, we show that these forces can be harnessed to promote the cellular uptake of specific receptors. The observation that two disordered domains with very different amino acid compositions, the low complexity domain of FUS and the RGG domain of LAF-1, have very similar effects on receptor uptake suggests that the phenomenon is not specific to the particular amino acid sequence of the disordered ligand. Further, by mutating specific residues within each sequence to modulate the strength of attractive interactions, we demonstrated that ligands can be engineered to tune the rate of receptor removal from the plasma membrane (Fig. 4). Notably, traditional bivalent monoclonal antibodies (IgG) were largely excluded from endocytic sites, and had minimal impact on receptor internalization (Fig. 2 and *SI Appendix*, Fig. S3), in line with literature reports, which suggest that the steric bulk of antibodies, in combination with their propensity for overly strong receptor clustering, makes them inappropriate for manipulating endocytic rates (20).

While moderate concentrations of attractive ligands promoted uptake of receptors, our results also revealed that excessive binding of disordered ligands to the cell surface inhibited uptake. This biphasic effect emerged simultaneously with the presence of rigid, plaque-like condensates of receptors and ligands on the cell surface, suggesting that productive endocytosis requires ligand-receptor condensates to remain flexible. These findings are consistent with the emerging view that maintaining a flexible state is essential for the function of biomolecular condensates in cells (42, 43). Moreover, the endocytic machinery requires structural plasticity, suggesting that overly stable or rigid assemblies can stall the process (44). Thus, while attractive interactions between ligand-receptor complexes can drive receptor uptake, their excessive condensation appears to create a physical barrier that inhibits membrane dynamics. Specifically, we speculate that moderate condensation generates compressive stresses that favor membrane bending, whereas overly dense assemblies may increase membrane rigidity and raise the energetic cost of membrane remodeling.

In contrast, repulsive interactions among ligands reduced recruitment of receptor-ligand complexes to endocytic sites (Fig. 7). This effect is analogous to the reduction in endocytic uptake of bulky glycosylated proteins, which have been shown to generate steric pressure in opposition to endocytosis (18, 45). By tethering repulsive disordered domains to our model receptor, we effectively created a steric barrier to endocytic uptake. These results suggest that ligands can be designed to retain specific receptors on the cell surface using steric exclusion from endocytic sites.

Collectively, our results establish a set of design principles for engineering disordered ligands that can control receptor internalization by endocytosis. By optimizing the amino acid composition and sequence features of intrinsically disordered proteins, it is possible to exert both positive and negative control over receptor uptake. The use of disordered ligands to enhance receptor uptake could be particularly valuable for therapeutic applications, such as enhancing the efficacy of receptor-degrading chimeras such as lysosome-targeting chimeras against oncogenic receptors such as receptor tyrosine kinases, where the efficiency of internalization is often a major limiting factor (46). Conversely, steric repulsion could be leveraged to stabilize proteins with short lifetimes at the plasma membrane, such as cystic fibrosis transmembrane conductance regulator in cystic fibrosis, glucose transporters in metabolic syndrome, and apoptotic receptors in cancer cells. Moving forward, further development of this tool will optimize ligand properties such as molecular weight and affinity to ensure efficient and specific control of the lifetime of receptors on the plasma membrane.

## Materials and methods

### Reagents

Tris-hydrochloride (Tris-HCl), 4-(2-hydroxyethyl)-1-piperazineethanesulfonic acid (HEPES), NaCl, KCl, isopropyl-β-D-thiogalactopyranoside (IPTG), β-mercaptoethanol, Triton X-100, Sodium bicarbonate, sodium tetraborate, phenylmethylsulfonyl fluoride (PMSF), Ethylene diamine tetraacetic acid (EDTA), Dithiothreitol (DTT), imidazole, glycerol, Poly-L-Lysine (PLL), Hydrocortisone, neutravidin, and Texas Red-DHPE were obtained from Sigma-Aldrich. POPC, DGS-NTA-Ni, and Biotinyl Cap PE were purchased from Avanti Polar Lipids. Amine-reactive PEG (mPEG-succinimidyl valerate MW 5000) and PEG-biotin (Biotin-PEG SVA, MW 5000) were purchased from Laysan Bio.

### Plasmids

Plasmid for purifying GFP-FUS LC (residues 2 to 214) was generously provided by the laboratory of Steven Mcknight at the University of Texas Southwestern Medical center(34). Plasmid for purifying GFP-FUS LC 1-163, GFP-FUS LC 1-163 StoQ, GFP-FUS LC 1-163 QtoS, GFP-RGG, GFP-KGG, and GFP-AP180CTD were custom-designed and constructed by GenScript (GenScript).

### Cell culture and Transfection

Human-derived SUM159 cells gene-edited to add HaloTag to both alleles AP-2 σ2 were a gift from T. Kirchhaus (32). Cells were grown in 1:1 Dulbecco’s Modified Eagle Medium (DMEM) high glucose: Nutrition mixture (F-12 HAM’s) (Hyclone, Cytiva) supplemented with 5 % fetal bovine serum (GeminiBio), Penicillin/Sterptomycin/L-glutamine (Cytiva), 1 μg mL−1 Hydrocortisone (Sigma-Aldrich), 5 μg mL−1 Insulin (Gibco) and 10 mM HEPES. Cells were incubated at 37 **°**C with 5% CO2. Cells were seeded onto acid-washed coverslips for 24 h before transfection with 1 μg of plasmid DNA using 3 μL Fugene HD transfection reagent (Promega). HaloTagged AP-2 σ2 was visualized by adding the JF646- HaloTag ligand (Promega). Ligand (60 nM) was added to cells and incubated at 37 **°**C for 10 min. Cells were washed with fresh medium and imaged immediately.

### Fluorescence microscopy

Live cell images were collected on an Olympus spinning disk confocal microscope, which featured a Yokogawa CSU-W1 SoRa confocal scanner unit and an Olympus IX83 microscope equipped with a 100xPlan-Aporchromat 1.5 NA oil-immersion objective. Fluorescence emission was captured using a Hamamatsu ORCA C13440-20CU CMOS camera. Lasers with excitation wavelengths of 488 nm for GFP, 561 nm for RFP, and 640 nm for JF646 were used.

Live cell FRAP experiments were performed using Olympus unit 405 nm laser on the spinning disk confocal microscope. A square region, 5 μm for each side, was photobleached on the plasma membrane of cells incubated with 1.25 μM GFP-FUS LC 1-163. The coverslip was heated to produce a sample temperature of 37 **°**C. Images were captured every 2 s for 2 min after photobleaching.

### Image analysis

Fluorescence images analyzed in ImageJ (http://rsbweb.nih.gov/ij/). Cell FRAP data were analyzed where fluorescence recovery over time was measured and then normalized to the maximum pre-bleach intensity. Recovery for endocytic sites was manually measured due to its high mobility of the endocytic structures on the plasma membrane. Recovery for the membrane and aggregated region was measured by drawing a rectangular window capturing the photobleached region.

Clathrin-coated structures were detected and tracked using cmeanalysis in MATLAB(33). Gaussian functions were fit to local intensity maxima in the AP-2 σ2 HaloTag labeled with JF646 (master channel), which marks the clathrin-coated structures. The Gaussian standard deviation was derived from the microscope’s physical parameters to estimate the point spread function. To ensure the accuracy of the fit, the Anderson-Darling test was applied to the residuals. The Gaussian amplitude, corresponding to the fluorescence intensity of each detected punctum, along with the punctum’s location, were recorded. For a punctum to qualify as a valid clathrin-coated structure, it had to be diffraction-limited and significantly brighter than the surrounding membrane, as described in previous work (33). For validated puncta in the master channel, a 2D Gaussian was then fit to the corresponding puncta in the receptor and/or ligand channels. A gaussian curve was fitted in the subordinate channels within a 3 σ pixel radius of the corresponding location in the master channel.

### Protein expression and purification

GFP-FUS 1-214, GFP-FUS 1-163, GFP-FUS 1-163 QtoS, GFP-FUS 1-163 StoQ, GFP-RGG, and GFP-KGG and GFP-AP180CTD constructs were transformed into *E. coli* BL21 (DE3) competent cells (NEB), which were grown at 30 °C to an optical density (OD) 600 of 0.8, and then cooled down to 16 °C. Protein expression was induced with 1 mM IPTG overnight at 16 °C with shaking at 200 rpm. The cells were pelleted from 1 L cultures by centrifugation at 4785 x g (5000 rpm in Beckman JLA8.1000) for 20min. The purification protocol was optimized for each construct as indicated (Table S1).

Each pellet was resuspended in 100 mL of the indicated lysis buffer to which 1% triton X-100 had been added by homogenization with a Dounce homogenizer followed by sonication (4 x 2000 J) on ice. The lysate was clarified by centrifugation at 29,535 x g (20,000 rpm in Beckman JA-25.50) for 30 min at 4 °C. The cell lysate was then incubated at the indicated temperature with 10 mL bed volume of Ni-NTA agarose (Qiagen) with gentle stirring at 60 rpm for 30 minutes, poured into a column, and washed with 10 column volumes of the indicated lysis buffer to which 0.2% triton X-100 had been added, followed by washing with 5 column volumes of detergent free lysis buffer. The protein was eluted with the indicated Ni-NTA Agarose elution buffer and concentrated with Amicon Ultra-15 centrifugal filters 10K MWCO (Thermo Scientific) down to 5-10 mL. Proteins that were adequately pure at this stage were further concentrated to >50 µM and dialyzed into the indicated storage buffer followed by centrifugation at 436,000 x g (100,000 rpm in Beckman rotor TLA 120.2) for 10 min at 4 °C to remove any aggregated protein. Single-use aliquots were flash-frozen using liquid nitrogen and stored at - 80 °C until the day of an experiment. Proteins that required additional purification were run on a Superdex 200 gel filtration column (Cytiva Hiload 26/600) with the indicated gel filtration buffer at 4 °C. After gel filtration the protein was centrifuged at 436,000 x g (100,000 rpm in Beckman rotor TLA 120.2) for 10 min at 4 °C to remove any aggregated protein. Single-use aliquots were flash-frozen using liquid nitrogen and stored at - 80 °C until the day of an experiment. Note, the group of proteins that were run on Ni-NTA agarose at 20 °C precipitated on ice, but were stable when run on a Superdex 200 column housed in a cold-box maintained at 4 °C (reported column temperature was 8-9 °C).

### GUV preparation

GUVs were made of 83% POPC, 15% Ni-NTA, and 2 mol % CAP biotin and an additional 0.1 mol% Texas Red DHPE lipids for visualization. GUVs were prepared following published protocol(47). Briefly, lipid mixtures dissolved in chloroform were spread into a film on indium tin oxide-coated glass slides (resistance, ∼8 to 12 W sq^-1^). And further dried in a vacuum desiccator for at least 2 hours to remove all of the solvent. Electroformation was performed at 55 °C glucose solution. 340 milliosmole glucose solution was used for making GUVs. The voltage was increased every 3 min from 50 to 1400 mV peak to peak for the first 30 min at a frequency of 10 Hz. The voltage was then held at 1400 mV peak to peak for 120 min and lastly was increased to 2200 mV peak to peak for the last 30 min during which the frequency was adjusted to 5 Hz. GUVs were stored in 4 °C and used within 3 days after electroformation.

### GUV tethering and imaging

GUVs were tethered to glass coverslips for imaging as previously described(38). Briefly, glass coverslips were passivated with a layer of biotinylated poly-L-lysine conjugated PEG chains (PLL-PEG). GUVs doped with 2 mol% biotinylated Cap PE were then tethered to the passivated surface using neutravidin.

PLL-PEG was synthesized as described previously with minor modifications as follows(38). Briefly, an amine-reactive mPEG succinimidyl valerate was conjugated to poly-L-lysine at a molar ratio of 1:5 PEG to PLL. The conjugation reactions take place in 50 mM sodium tetraborate (pH 8.5), and are allowed to react overnight at room temperature while continuously stirring. The final product is then buffer exchanged to PBS (pH 7.4) using 7000 MWCO Zeba spin desalting columns (Thermofisher) and stored at 4 °C.

Imaging wells consisted of 5 mm diameter holes in 0.8 mm thick silicon gaskets (Grace Bio-Labs). Gaskets were placed directly onto no.1.5 glass coverslip (VWR International), creating a temporary waterproof seal. Before well assembly, gaskets and coverslips were cleaned in 2% (v/v) Hellmanex III (Hellma Analytics) solution, rinsed thoroughly with water, and dried under a nitrogen stream. The GUV buffer used for dilution and rinsing was 25 mM HEPES, 150 mM NaCl pH 7.4 buffer. In each dry imaging well, 20 μL of PLL-PEG was added, incubated 20 min and then the wells were serially rinsed with the buffer by gently pipetting. Next, 6 μg of neutravidin dissolved in 25 mM HEPES and 150 mM NaCl (pH 7.4) was added to each sample well and allowed incubation for 10 min. Wells were rinsed with the buffer and then 10 μL of diluted GUVs was added to the well and allowed incubation for 10 min. GUVs were diluted in the buffer at a ratio of 1:12. Excess GUVs were then rinsed from the well using the buffer, and the sample was subsequently imaged using confocal fluorescence microscopy

Imaging experiments were performed using spinning disk confocal microscopy. Image stacks taken at fixed distances perpendicular to the membrane plane (0.5-μm steps) were acquired immediately after GUV tethering and again protein addition. At least 30 fields of views were randomly selected for each sample for further analysis before and after the addition of protein, respectively. Imaging proceeded 5 min after adding protein to achieve protein binding and reaching a relatively equilibrium state.

### Calibration of the brightness of GFP and RFP molecules

The number of molecules was estimated on the basis of fluorescence intensity using previously reported protocol(16). Briefly, picomolar concentrations of purified GFP or RFP proteins were added to imaging wells containing RCA-cleaned coverslips, following ultracentrifugation at 100,000 x g to remove aggregates. Images of sparse-diffraction limited puncta were acquired using laser power and exposure settings identical to those used in live-cell imaging experiments. Particle detection software (described above for cell imaging experiments) was used to detect puncta and measure the amplitude of the Gaussian function that best fit these puncta. To validate that these puncta represented single molecules, we confirmed single-step photobleaching behavior. The resulting Gaussian fluorescence amplitude was linearly scaled for exposure time and used to extrapolate the number of molecules in diffraction-limited endocytic structures.

### Protein labeling

The GFP antibody (sc-9996, Santa Cruz Biotechnology) used in this study was labeled with Atto594, an amine-reactive, NHS ester-functionalized fluorescent dye. The labeling reaction took place in phosphate buffered saline supplemented with 10 mM of sodium bicarbonate. A dye was added to the protein in 10-fold stoichiometric excess and allowed reaction for 1 h at room temperature, resulting in a labeling ratio of 0.5 dyes per protein. Unconjugated dye was separated using Zeba Dye and Biotin Removal Spin Columns.

### Statistical analysis

All experiments were repeated independently on different days at least three times, with similar results. ImageJ was used to analyze confocal images. Statistical analysis was carried out using a two-tailed Student’s *t*-test (unpaired, unequal variance). Error bars in graphs represent either standard error or standard deviation as stated in figure captions.

## Supplementary data

**Fig. S1.**
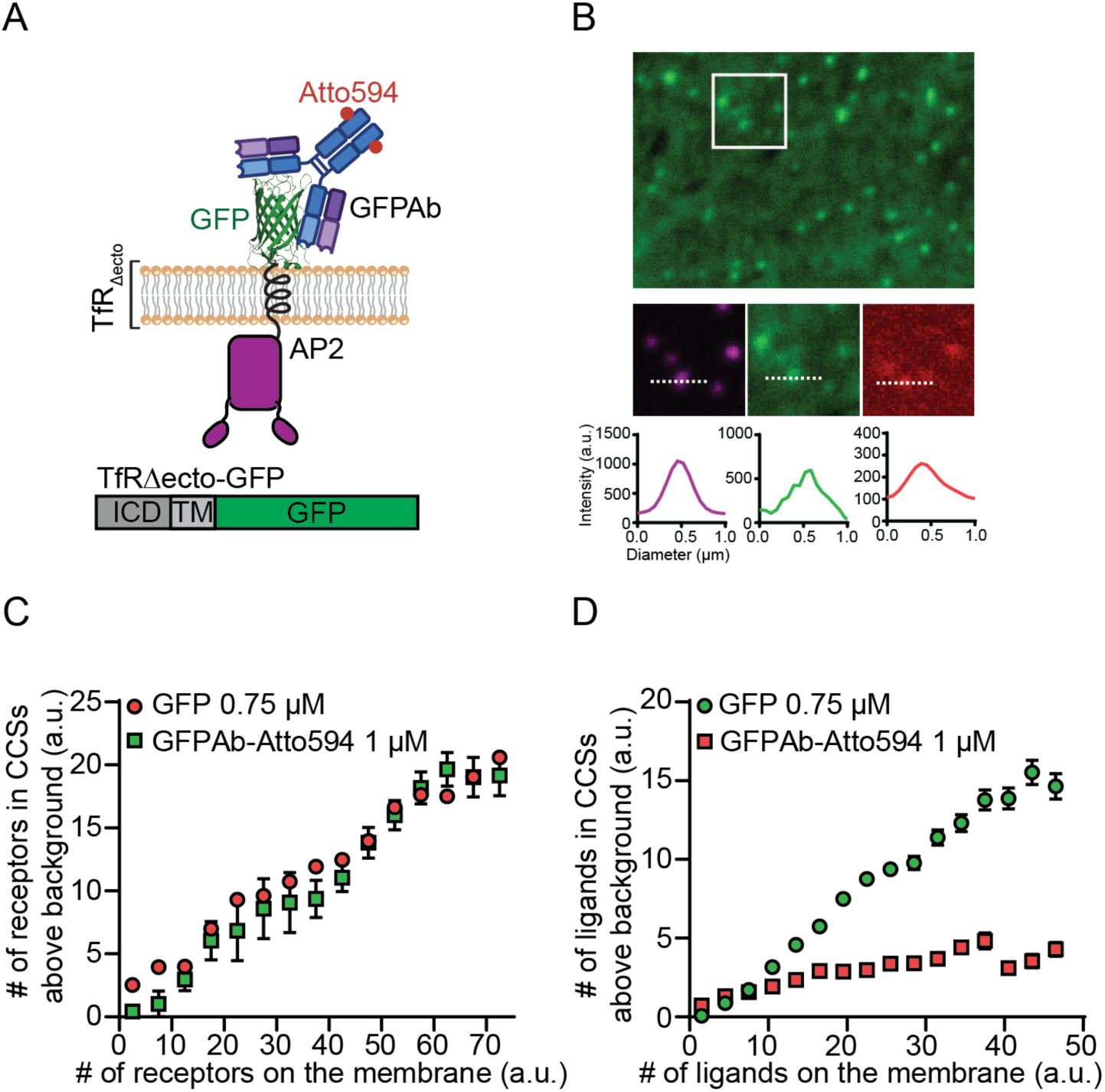
Monoclonal IgGs do not effectively promote receptor internalization. **A.** Schematic illustration of the transferrin receptor-based model receptor (TfRΔecto-GFP) bound by an Atto594-labeled anti-GFP monoclonal antibody (GFPAb). The model receptor consists of the intracellular and transmembrane domains of the TfR fused to an extracellular GFP domain. **B.** Spinning disk confocal microscopy images of plasma membranes of SUM159 cells transiently expressing the Transferrin receptor tagged with GFP. Endocytic sites were visualized using HaloTag-JF646-labeled AP2 (magenta). Cells were incubated with 1 µM Atto594-labeled anti-GFP monoclonal antibody (GFPAb-Atto594). The white box indicates the magnified inset, and graphs show intensity lines profiled across the dashed lines. **C-D**. The representative curves of the estimated absolute number of **(C)** receptors and **(D)** ligands within endocytic proteins localized in CCSs are shown against the estimated absolute number of receptors and ligands on the plasma membrane around each structure. Fluorescence intensities for 0.75 µM GFP and 1 µM GFPAb-Atto594 were divided by their respective single-molecule calibration values (*SI Appendix*, Fig. S6) to estimate the number of molecules. Error bars represent mean ± SE.

**Fig. S2.**
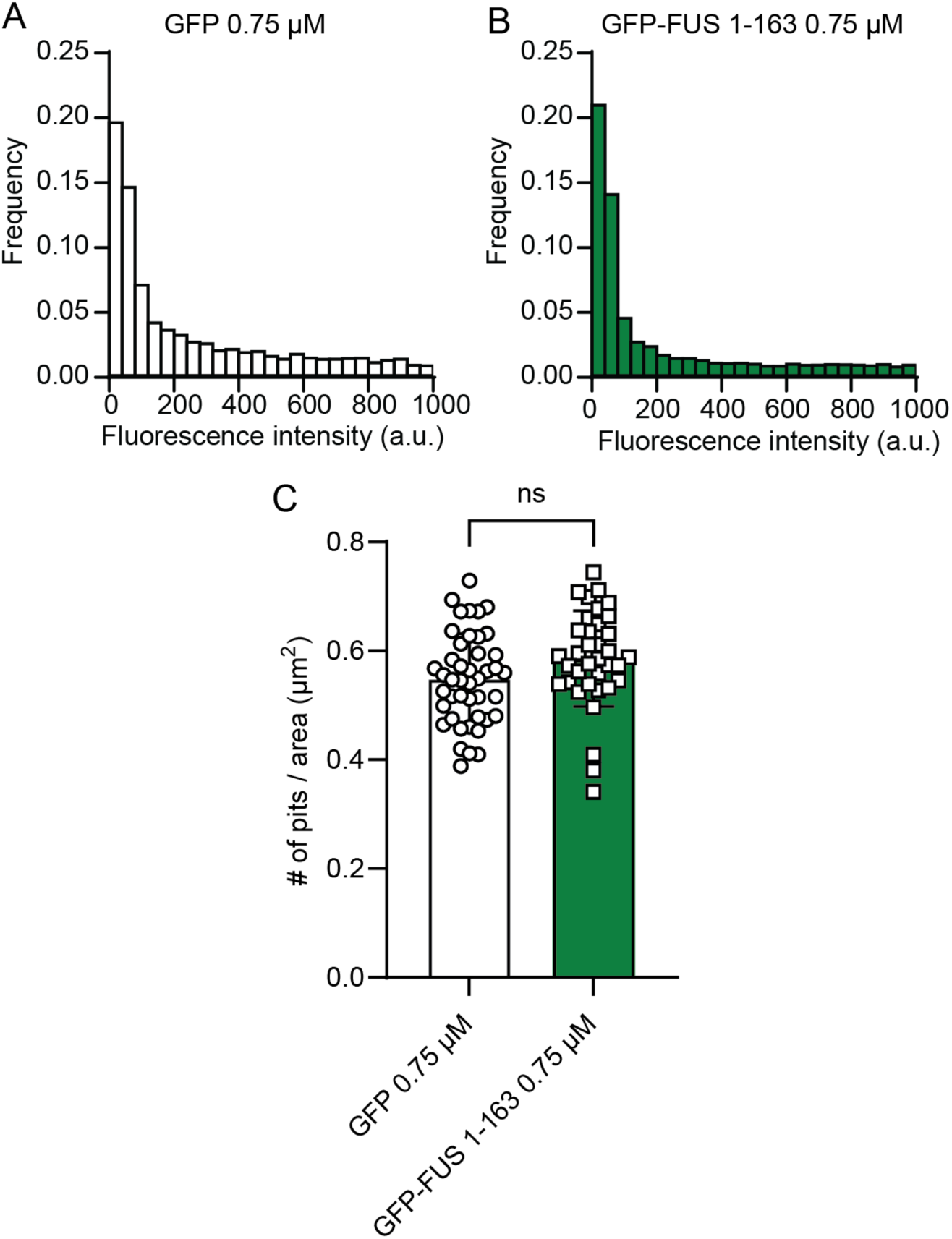
Comparison of AP2 fluorescence intensity between GFP and GFP-FUS 1-163. **A-B**. Histogram of the AP2 tracked in endocytic pits when cells were exposed to **(A)** 0.75 µM GFP and **(B)** 0.75 µM GFP-FUS 1-163. A total of 4,907 endocytic sites were detected from 44 cells incubated with 0.75 µM GFP, and 10,366 endocytic sites were detected from 54 cells incubated with 0.75 µM GFP-FUS 1-163. **(C)** Bar chart comparing the density of endocytic pits between GFP and GFP-FUS 1-163. Density was calculated as the number of detected AP2 divided by the cell surface area. n = 45 and 36 regions were analyzed for each plot, respectively. Error bars represent mean ± SD. Statistical significance was tested using an unpaired, two-tailed Student’s *t* test (n.s. means no significant difference).

**Fig. S3.**
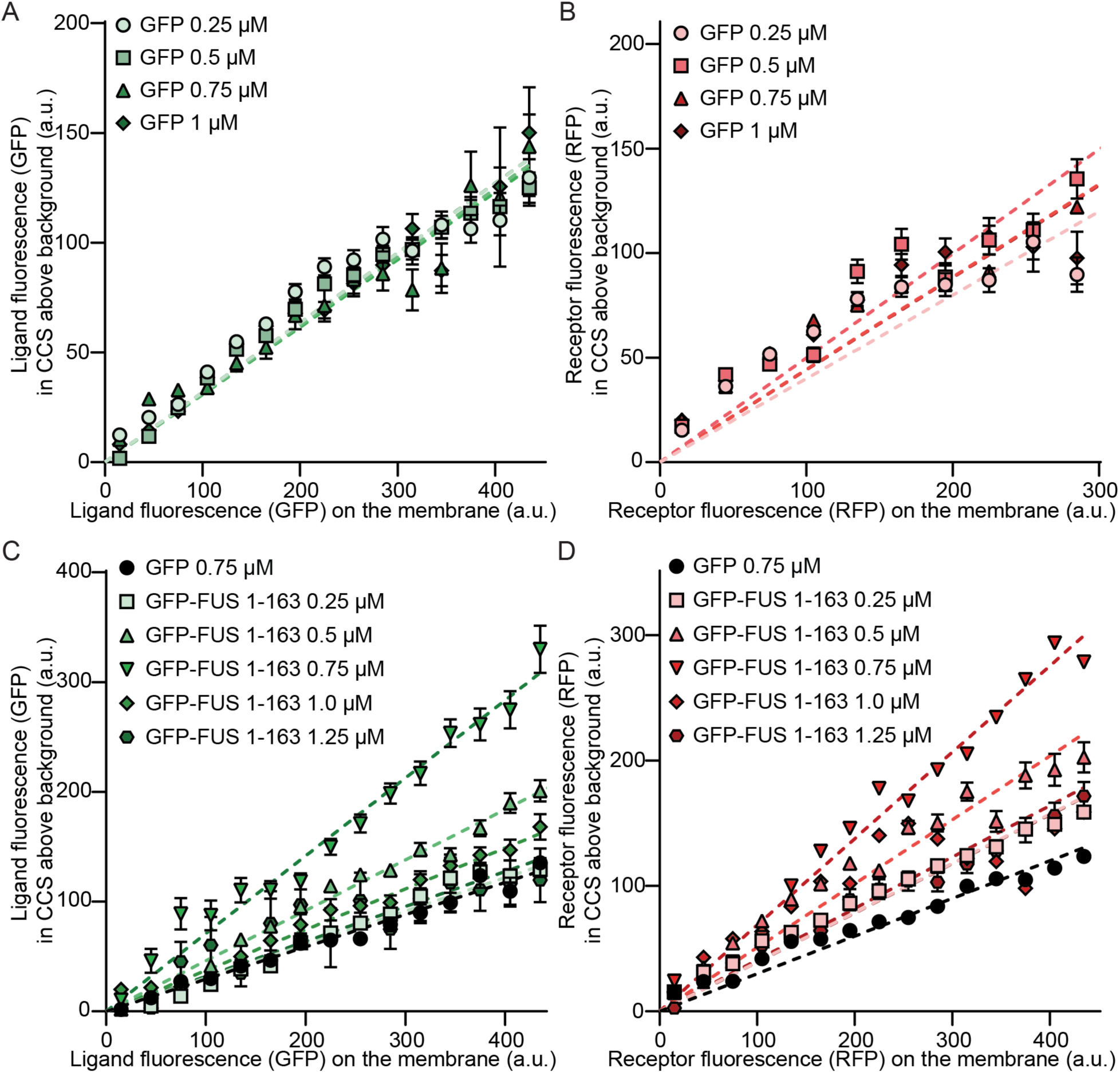
CMEanalysis curves underlying slope calculations for GFP and GFP-FUS 1-163 conditions. **A-B.** CMEanalysis curves for the GFP ligand at the indicated concentrations. The representative curves of relative number of **(A)** ligands and **(B)** receptors within endocytic proteins localized in clathrin-coated structures are shown against the relative concentrations of model ligands and receptors on the plasma membrane around each structure. A total of 6,717 endocytic sites were detected from 51 cells incubated with 0.25 µM GFP, 11,889 endocytic sites were detected from 51 cells incubated with 0.5 µM GFP, 4,907 endocytic sites were detected from 44 cells incubated with 0.75 µM GFP, and 16,065 endocytic sites were detected from 55 cells incubated with 1.0 µM GFP. **C-D**. Representative curves for GFP-FUS 1-163 across concentrations for **(C)** ligands and **(D)** receptors. A total of 4,907 endocytic sites were detected from 44 cells incubated with 0.75 µM GFP, 8,352 endocytic sites were detected from 47 cells incubated with 0.25 µM GFP-FUS 1-163, 6,979 endocytic sites were detected from 41 cells incubated with 0.5 µM GFP-FUS 1-163, 10,366 endocytic sites were detected from 54 cells incubated with 0.75 µM GFP-FUS 1-163, 9,819 endocytic sites were detected from 53 cells incubated with 1.0 µM GFP-FUS 1-163, and 12,816 endocytic sites were detected from 54 cells incubated with 1.25 µM GFP-FUS 1-163. Dashed lines indicate linear fits used to extract slope values, revealing optimal recruitment at intermediate GFP-FUS 1-163 concentrations and reduced recruitment at higher concentrations. Error bars represent mean ± SE.

**Fig. S4.**
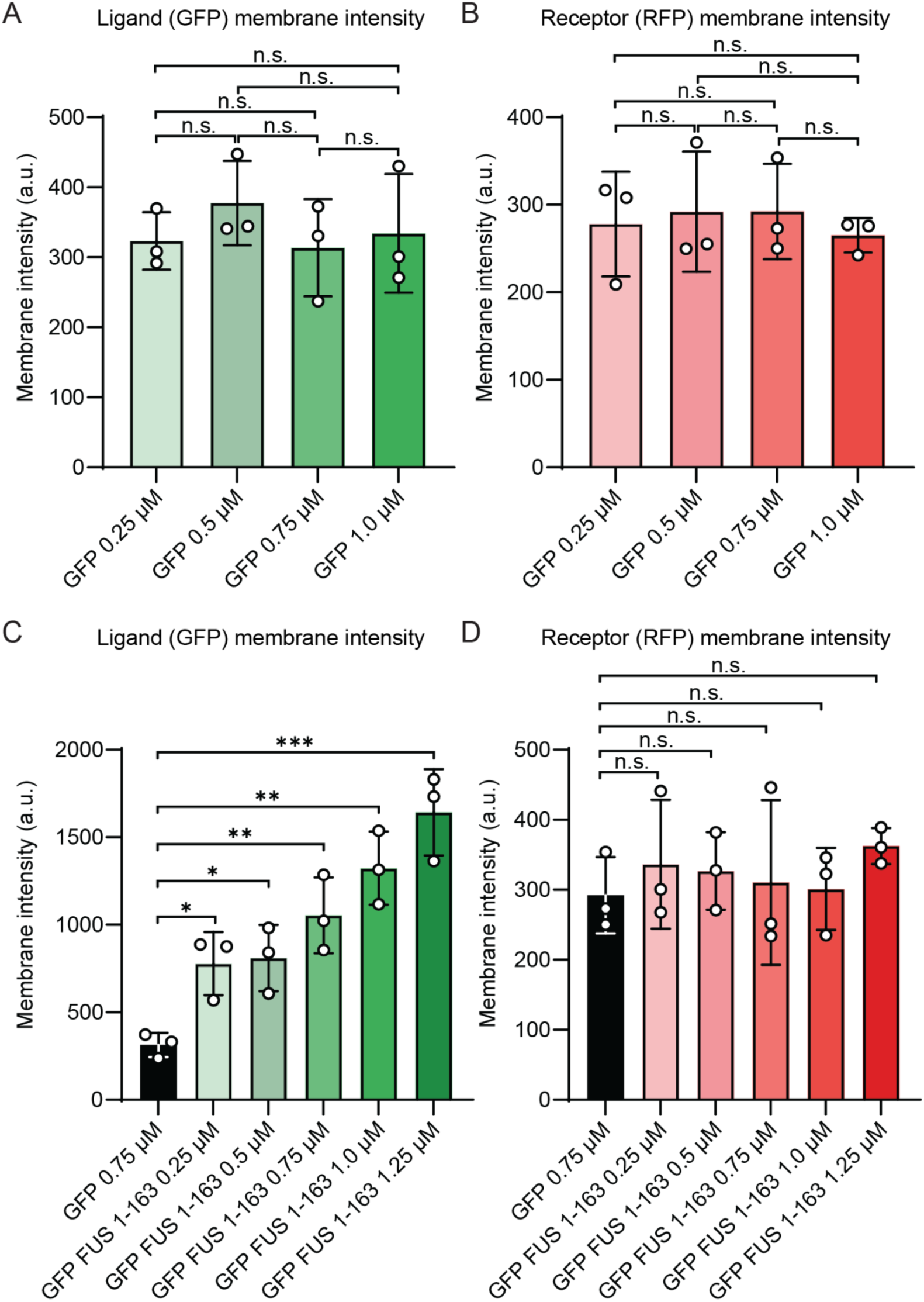
Ligand controls membrane intensity for GFP-FUS 1-163 but not for GFP ligand, while receptor membrane intensity remains unchanged. **A-B**. Bar chart comparison of **(A)** GFP and **(B)** RFP membrane intensity in SUM159 AP-Halo cells incubated with soluble GFP at the indicated concentrations. Neither the ligand (GFP) nor the model receptor (RFP) showed dose-dependent changes in membrane intensity under GFP ligands. **C-D**. Bar chart comparison of **(C)** GFP and **(D)** RFP membrane intensity in cells incubated with GFP-FUS 1-163. GFP membrane intensity increased with GFP-FUS 1-163 concentration, consistent with increased ligand concentration on the membrane, whereas RFP membrane intensity remained unchanged across conditions. Error bars show mean ± SD (*P < 0.05, **P < 0.01, ***P < 0.001, n.s. means no significant difference).

**Fig. S5.**
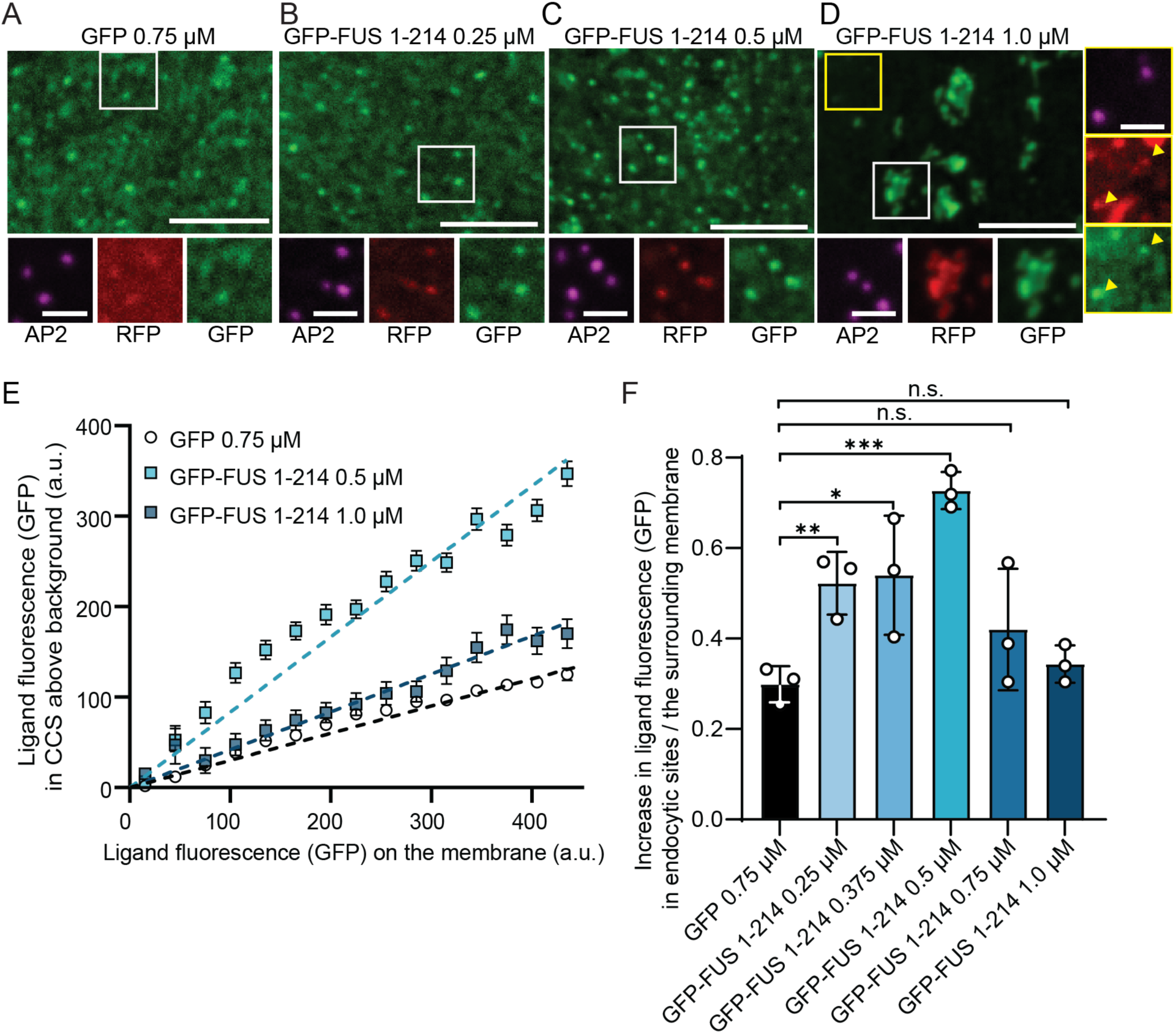
Increased ligand length promotes efficient endocytic uptake. **A-D**. Spinning disk confocal microscopy images of the plasma membrane of SUM159 cells transiently expressing the model receptor, visualized with HaloTag-JF646-labeled AP2 (magenta). Cells were incubated with **(A)** 0.75 µM GFP, **(B)** 0.25 µM GFP-FUS 1-214, **(C)** 0.5 µM GFP-FUS 1-214, or **(D)** 1 µM GFP-FUS 1-214. The white boxes indicate the magnified insets and yellow boxes in (D) indicate magnified insets showing dim puncta corresponding to endocytic sites. The scale bars are 5 µm (main panels) and 1 µm (insets). **E**. The relative number of model receptors within endocytic proteins localized in clathrin-coated structures is shown against the relative concentration of fusion proteins on the plasma membrane around each structure. A total of 4,907 endocytic sites were detected from 44 cells incubated with GFP, 16,037 endocytic sites were detected from 57 cells incubated with 0.5 µM GFP-FUS 1-214, and 14,065 endocytic sites were measured from 56 cells incubated with 1.0 µM GFP-FUS 1-214. Error bars represent mean ± SE. **E**. Slope analysis for various concentrations of GFP-FUS 1-214. A total of 13,618 endocytic sites were detected from 138 cells incubated with GFP, 46,310 endocytic sites were detected from 172 cells incubated with 0.25 µM GFP-FUS 1-214, 31,349 endocytic sites were measured from 202 cells incubated with 0.375 µM GFP-FUS 1-214, 31,365 endocytic sites were detected from 177 cells incubated with 0.5 µM GFP-FUS 1-214, 34,440 endocytic sites were detected from 195 cells incubated with 0.75 µM GFP-FUS 1-214, and 37,715 endocytic sites were detected from 165 cells incubated with 1.0 µM GFP-FUS 1-214. Error bars represent mean ± SD from three independent experiments. Statistical significance was tested using an unpaired, two-tailed Student’s *t* test (*P<0.05, **P < 0.01, ***P < 0.001, n.s. means no significant difference).

**Fig. S6.**
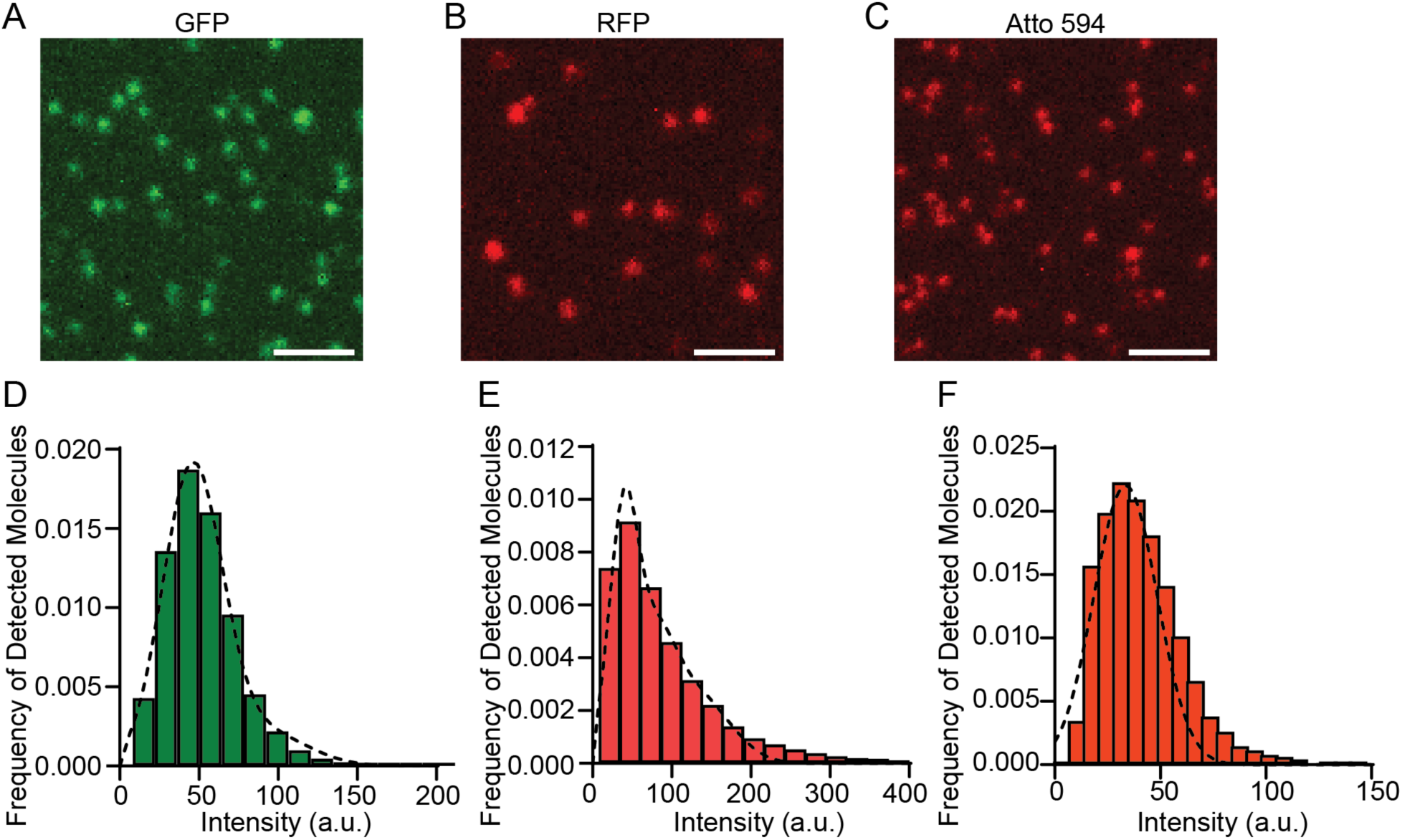
Measuring the intensity of individual fluorescent dye. **A-C**. Representative image of **(A)** GFP, **(B)** RFP, **(C)** ENTH-Atto594 proteins adhered passively to a coverslip surface. Samples were diluted to picomolar concentrations to obtain sparse, diffraction-limited puncta on analysis. The scale bar is 2 µm.**D-F**. Histogram showing the frequency distribution of fluorescence intensities for detected single **(C)** GFP, **(E)** RFP, and **(F)** ENTH-Atto594 molecules. The dashed curves represent gaussian fits to the distributions. The peak value of the gaussian fit was defined as the calibrated brightness of a single molecule and used to estimate the receptor-ligand binding in live cells.

**Fig. S7.**
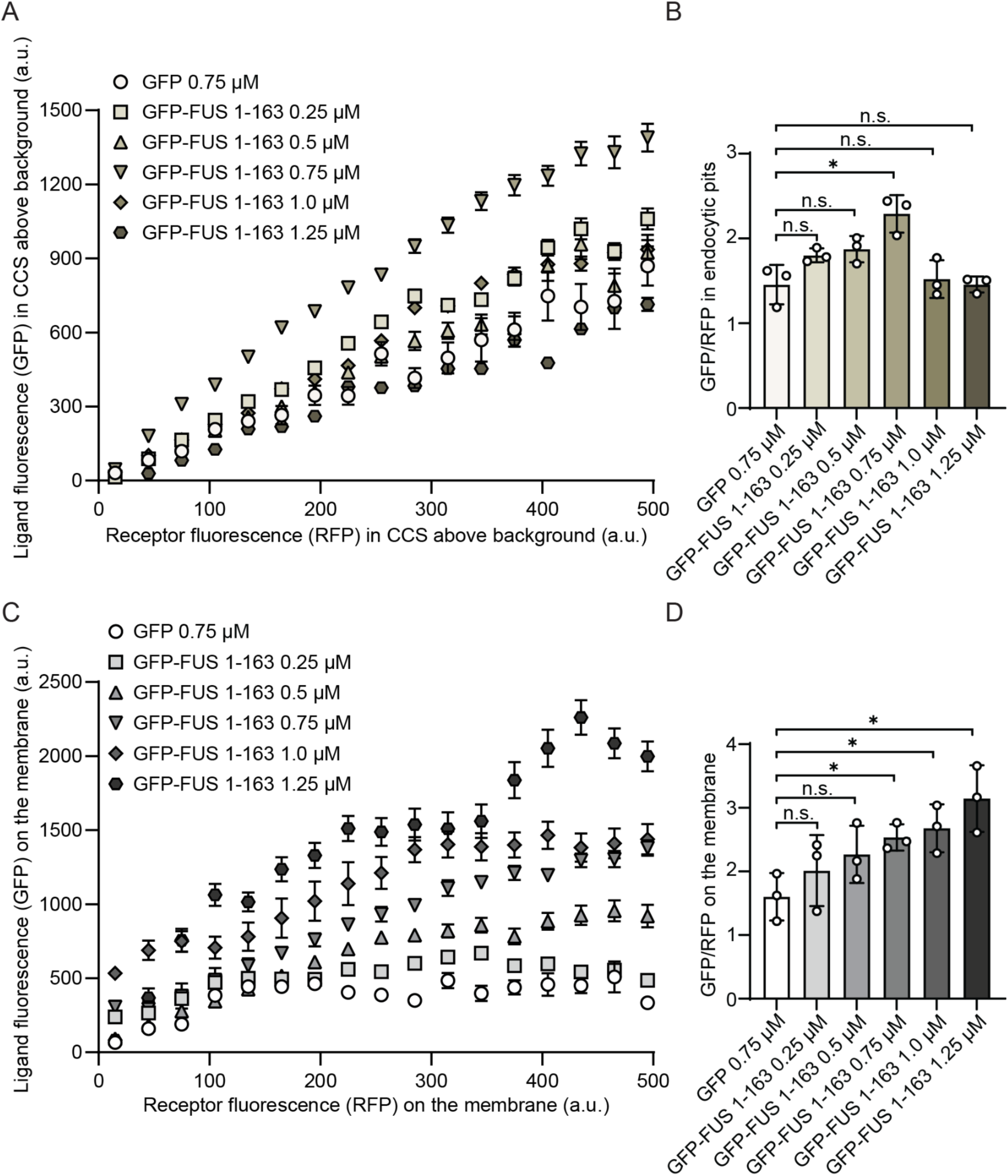
Relative ligand abundance at endocytic pits and on the membrane across GFP-FUS 1-163 concentrations. **(A)** Relationship between ligand and receptor enrichment at endocytic sites. Representative curves show GFP fluorescence in endocytic sites is plotted as a function of RFP fluorescence in endocytic sites for GFP and GFP-FUS 1-163 at the indicated concentrations. A total of 4,907 endocytic sites were detected from 44 cells incubated with 0.75 µM GFP, 8,352 endocytic sites were detected from 47 cells incubated with 0.25 µM GFP-FUS 1-163, 6,979 endocytic sites were detected from 41 cells incubated with 0.5 µM GFP-FUS 1-163, 10,366 endocytic sites were detected from 54 cells incubated with 0.75 µM GFP-FUS 1-163, 9,819 endocytic sites were detected from 53 cells incubated with 1.0 µM GFP-FUS 1-163, and 12,816 endocytic sites were detected from 54 cells incubated with 1.25 µM GFP-FUS 1-163. **(B)** Ratio of ligand to receptor signal within endocytic sites across concentrations, summarizing the relative ligand enrichment per receptor in endocytic sites. **(C)** GFP fluorescence on the surrounding plasma membrane is plotted as a function of RFP fluorescence on the membrane for GFP and GFP-FUS 1-163 across concentrations, showing increased membrane ligand signal with increasing GFP-FUS 1-163 dose. **(D)** Ratio of ligand to receptor signal on the membrane across conditions. Error bars in (A) and (C) represent mean ± SE. Error bars in (B) and (D) denote mean ± SD. Statistical significance was tested using an unpaired, two-tailed Student’s *t* test (*P<0.05, n.s. means no significant difference).

**Table S1.**
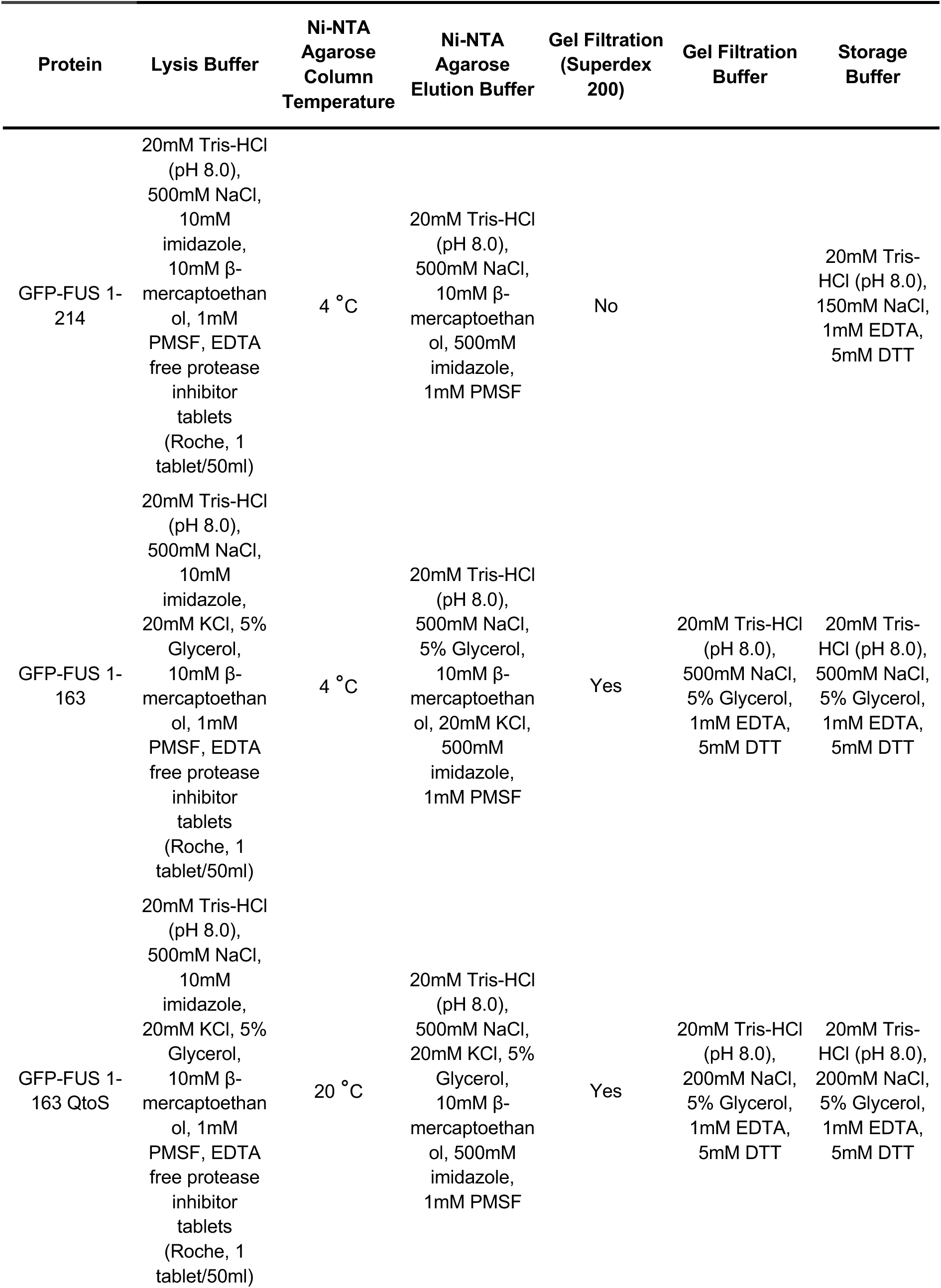
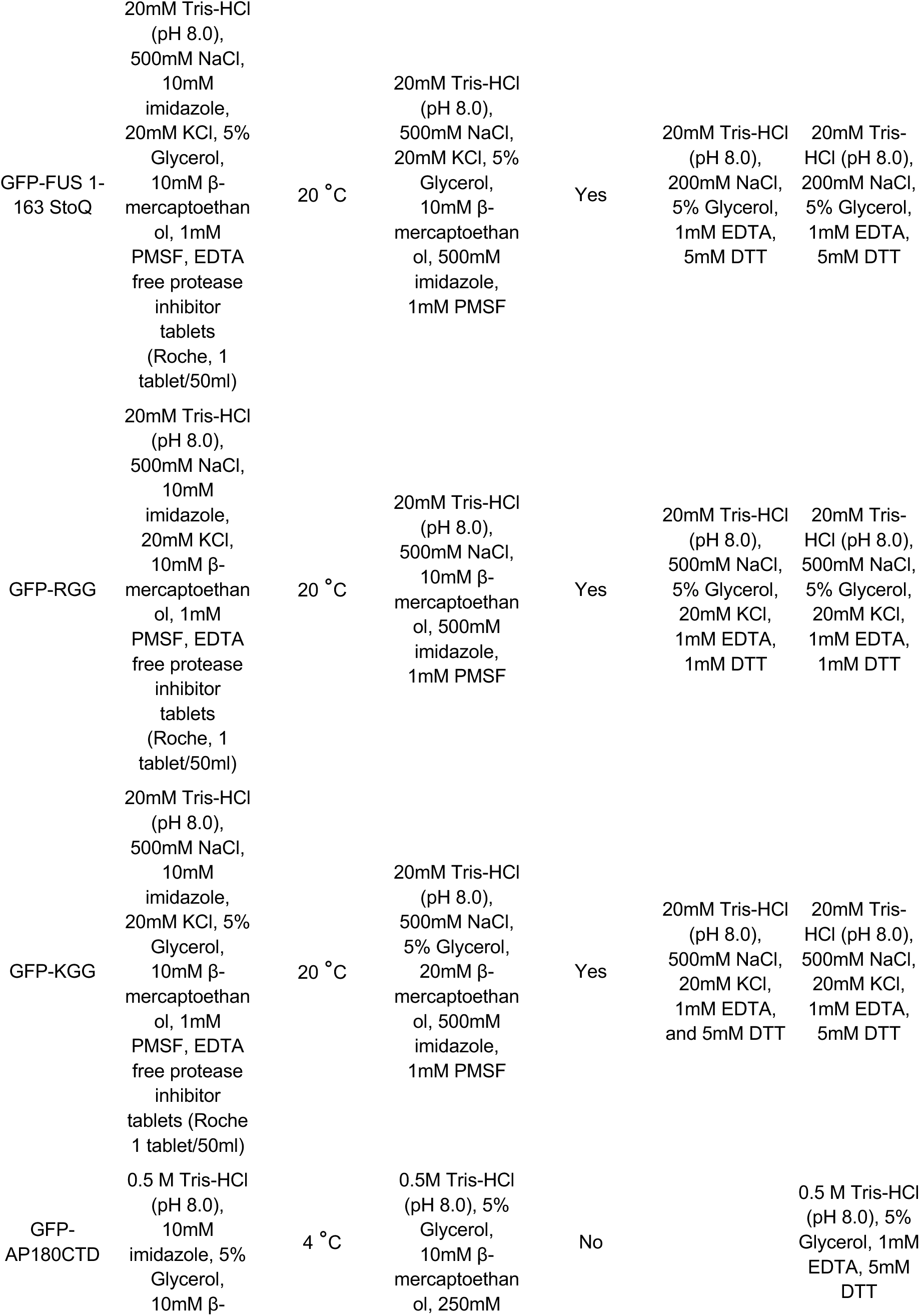
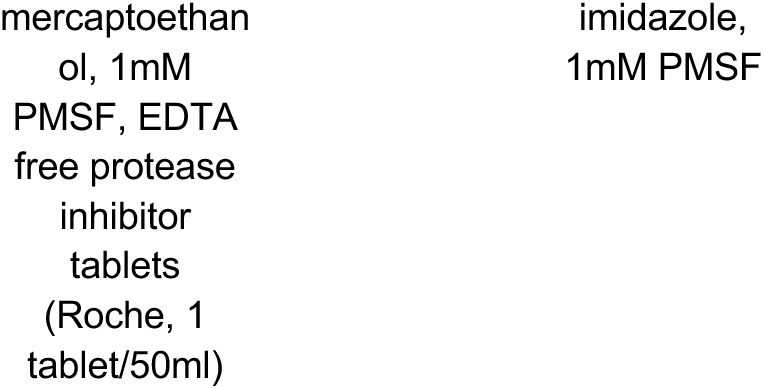
Summary of protein purification buffer compositions for different proteins.

## Notes

### Competing Interest Statement

The authors have declared no competing interest.

### Funding

- National Institute of General Medical Sciences: R35GM139531

